# DNA Polymerases in Precise and Predictable CRISPR/Cas9-mediated Chromosomal Rearrangements

**DOI:** 10.1101/2023.02.13.528245

**Authors:** Mohammadreza M. Mehryar, Xin Shi, Jingwei Li, Qiang Wu

**Author notes:** Joint Authors (Mohammadreza M. Mehryar, Xin Shi).

## Abstract

Recent studies have shown that Cas9-mediated CRISPR gene editing outcomes at junctions of chromosomal rearrangements are precise and predictable; however, the underlying mechanisms are poorly understood due to lack of suitable assay system and analysis tool. Here we developed a customized computer program to take account of staggered Cas9 cleavage and to rapidly process large volumes of junctional sequencing reads from chromosomal rearrangements, including DNA-fragment inversions, duplications, and deletions. We also established a sensitive assay system using *HPRT1* and *DCK* as reporters for cell growth during DNA-fragment editing by Cas9 with dual sgRNAs and found prominent large resections at junctions of chromosomal rearrangements. In addition, we found that knockdown of *PolQ* (encoding Polθ) results in increased large resections but decreased small deletions. We also found that the mechanisms for generating small deletions of 1bp and >1bp during DNA-fragment editing are different with regards to their opposite dependencies on Polθ and Polλ. Specifically, Polθ suppresses 1bp deletions but promotes >1bp deletions, whereas Polλ promotes 1bp deletions but suppresses >1bp deletions. Finally, we found that Polλ is the main DNA polymerase responsible for fill-in of the 5’ overhangs of staggered Cas9 cleavage ends. These findings contribute to our understanding of the molecular mechanisms of CRISPR/Cas9-mediated DNA-fragment editing and have important implications for controllable, precise, and predictable gene editing.

## INTRODUCTION

CRISPR gene editing outcomes result from cellular ligations of double strand break (DSB) ends after Cas9 cleavages. This occurs either via homologous recombination (HR) during the S and G2 cell cycle phases or via non-homologous end joining (NHEJ) throughout all four phases of the cell cycle. The former results in precise modifications while the latter is associated with random indels (1-4). However, a major outstanding issue, particular with regards to the NHEJ process, is our incomplete understanding of the underlying mechanisms, including the identity of the cellular DNA polymerases that are involved in repairing the DSB ends following Cas9 cleavages (5-9).

An excellent model system to obtain a better understanding of CRISPR gene editing is the use of dual sgRNAs to study Cas9-mediated chromosomal rearrangements (4,10-12). In particular, Cas9 programmed with dual sgRNAs can result in chromosomal rearrangements including DNA-fragment deletions, inversions, and duplications (Supplementary Figure S1) (13-16). Details of this mechanism can thus also inform processes associated with normal chromosomal rearrangements, which are known to promote genome instability in cancers or generate immune diversity during development and allelic diversity during meiosis (17-19).

The advantage of using Cas9 with dual sgRNAs over single sgRNAs is that the repair outcomes of chromosomal rearrangements cannot be recut whereas there is repeated cutting and re-ligation for Cas9 with single sgRNAs (20,21). Consequently, repair outcomes of Cas9-mediated nucleotide insertions at ligation junctions of chromosomal rearrangements are more precise and predictable than those at editing sites with single sgRNAs (20,22-30). These precise insertions of predictable nucleotides at editing sites are thought to be resulted from fill-in and ligation of staggered DSB ends of Cas9 cleavages. However, the underlying DNA polymerase(s) remains unknown. Here we systematically analyzed the role of DNA polymerases in CRISPR/Cas9-mediated chromosomal rearrangements and found prominent roles of Polλ and Polθ in processing DSB ends during DNA-fragment editing yet with unexpected specificities.

## MATERIALS AND METHODS

### Cell culture

The human endometrial carcinoma HEC-1-B cells were cultured in the modified Eagle’s medium (MEM) supplemented with 10% fetal bovine serum (FBS) and 1% penicillin-streptomycin at 37°C in a 5% (v/v) CO_2_ incubator.

### SgRNA design and plasmid construction

We designed sgRNA sequences using CRISPOR, most of which were located at DNase I hypersensitive sites. The plasmid construction was performed as previously described (24). In brief, the pGL3-U6-sgRNA-PGK-puro was linearized with BsaI (NEB) at 37°C for 1.5h. The resulting plasmid backbone of the linearized vector was run on the 0.8% agarose gel and purified by Monarch DNA Gel Extraction Kit (NEB). The oligos for the inserted sgRNA targeting sequences were synthesized with two overhangs compatible with the linearized vector and complementary to each other. After annealing, the duplexes were ligated with the purified vector with T4 DNA ligase (NEB). The ligation products were transformed into DH5α bacteria for amplification. All plasmids were confirmed by Sanger sequencing.

### DNA polymerase knockdown

Knockdown experiments were performed as previously described (24,27).Briefly, we designed two sgRNAs for each polymerase, both targeting coding regions to achieve an efficient knockdown. HEC-1-B cells were plated in 6-well plates with around 30-40% cell confluence one day before transfection. When cells reached more than 80% confluence, they were co-transfected with Cas9 and two sgRNAs plasmids using Lipofectamine 3000 (Thermo Fisher) according to the manufacturer’s instructions. After 12 hours of culturing with 5% FBS, the culture medium was changed back to the normal condition with 10% FBS. Culturing continued for an additional 12 to 24 hours, then cell growth was assayed with the *HPRT*1 and *DCK* reporter systems.

#### *HPRT1* and *DCK* assay systems

We used the two reporter systems of *HPRT1* or *DCK* to detect large resections of the flanking exons induced by intronic targeting sites by the CRISPR/Cas9 system with dual sgRNAs. For the *HPRT1* assay system, cells with functional *HPRT1* are very sensitive to the 6-TG (6-thioguanine) chemical, and convert it into toxic thioguanosine monophosphate. By contrast, cells with deficient or non-functional *HPRT1* are resistant to this lethal drug and can survive. Cells without *DCK*, a housekeeping gene that plays an important role in DNA synthesis, are not able to accomplish DNA synthesis and will end with cell apoptosis. We designed dual sgRNAs 70-100 bp away from the splicing site within the intron 2 and intron 4 of *HPRT1* and *DCK*, respectively. If there are large resections into the flanking exons of *HPRT1* or *DCK* induced by Cas9 with dual sgRNAs, cells will survive in the *HPRT1* or die in the *DCK* assay systems.

For the *HPRT1* cell growth assay, we plated HEC-1-B cells on the 6-well plate with a cell confluence of 30-40% one day before transfection. The number of cells plated in each well were kept consistent. When cell confluence reached 80%, we transfected the cells with plasmids targeting different polymerases to obtain knockdown cell populations. Two days later, cells were transfected again with sgRNAs targeting intron 2 of *HPRT1* in the low serum medium. The medium was changed back to normal serum and continued culturing for one more day. Finally, we selected cells with 6-TG at a concentration of 10 μg/ml for 7 consecutive days. The cells were collected to count the numbers on Day1, Day2, Day4, Day6, and Day7. For the *DCK* cell growth assay, the procedures were similar but without the use of 6-TG, and cells were collected on Day1, Day2, Day3, Day4, and Day5.

### Genomic DNA extraction

We extracted genomic DNA from transfected cells to obtain purified DNA templates for further analysis. Briefly, DPBS was used to collect cells when cell confluence reached 70%-80%. After centrifugation and discarding the supernatant, the cell pellets were resuspended in the lysis buffer (200 mM NaCl, 10 mM Tris-HCl (pH 7.4), 2 mM EDTA (pH 8.0) and 0.2% (wt/vol) SDS) and incubated at 37°C with 750 rpm overnight. The genomic DNA was precipitated with 0.7x volume of isopropyl alcohol after centrifuging at a high speed of 14,000 g for 0.5h. Finally, the pellet was washed with 80% ethanol and DNA was dissolved with TE. The genomic DNA can be stored at -20°C for at least half a year.

### Preparation of junctional amplicon libraries

Chromosomal rearrangements including fragment deletion, inversion, and duplication can be induced by Cas9 with two sgRNAs. We used high throughput sequencing to assay various junctional repair outcomes of different chromosomal rearrangements. Considering the limitation of the read length, the primers used here were all near the junctional site and the length of the final amplified products was less than 290 bp. The experiments were performed as previously described with modifications (20). Briefly, the primers were designed to be compatible with the Illumina sequencing platform. The PCR conditions were as follows: initial denaturation at 95°C for 3 min, 30 cycles of denaturation at 95°C for 30 sec, annealing at 60°C for 15 sec, and extension at 72°C for 30 sec, followed by a final extension at 72°C for 3 min. The PCR products were purified with the High-Pure PCR Product Purification kit (Roche) and then sequenced by the X ten platform.

### Multiplex high throughput sequencing

For assessing the junctional repair outcomes of each chromosomal rearrangement upon perturbing DNA polymerases, we constructed libraries using Illumina P5/P7 primers with unique barcodes and indexes. For cost-effective sequencing, we constructed libraries for the same experiment but different replicates with the same index and barcode, but split samples of replicates into different lanes for efficient sequencing. After library construction, we quantified libraries with Qubit dsDNA HS assay and pooled samples of different experiments with equal mole for efficient sequencing. We performed each polymerase knockdown experiment with three replicates. In total, we constructed 829 libraries for high-throughput sequencing.

### RNA extraction and RT-PCR

We used the TRIzol Reagents (Invitrogen) to obtain the total mRNA for the RT-PCR test. In detail, we used 1ml TRIzol reagent for each well of six-well plates with cell confluence of more than 80%. After homogenization, the samples were incubated for 5 min at room temperature to complete the dissociation of nucleoprotein complexes. Then 200 μl of chloroform was added to the samples, which were then shaken continuously vigorously for 15 sec. After shaking, samples were left at room temperature for 5 min, then spun at 12,000 g for 15 min at 4°C. After centrifugation, RNA was precipitated with 500 μl isopropyl alcohol. Finally, the pellets were washed with 75% ethanol twice and dissolved with RNase-free water. The RNA can be stored at -20°C for up to a year. For RT-PCR, we used HiScript III RT SuperMix (Vazyme) for reverse transcription according to the manufacturer’s instructions followed by PCR with targeting primers. Primer sequences are listed in Supplementary Table S1.

### Simultaneous sequencing of deletion and inversion junctions

LAM-HTGTS (Linear Amplification Mediated High-throughput Genomic Translocations Sequencing) was first introduced to detect translocations (31). We used this method with a few modifications to assay large resections at junctional sites of chromosomal rearrangements induced by CRISPR/Cas9 systems with dual sgRNAs. Briefly, HEC-1-B cells were plated on the 6-well plate with a cell confluence of around 30% one day before transfection. When cell confluence reached 70%, we added fresh medium with 5% FBS and performed transfection with Lipofectamine 3000 (Thermo Fisher) according to the manufacturer’s instructions. The medium was changed back to the normal medium 24h later and continued culturing for another day to obtain total genomic DNA. We dissolved genomic DNA at a final concentration of 250 ng/μl for sonication. The sonication conditions were 8 trains of 30 sec ON and 90 sec OFF with low intensity. After sonication, the fragmented DNA was analyzed on 1.5% agarose gel and the ideal size should be 400-600 bp.

To acquire junctional repair outcomes of inversion and deletion simultaneously, we used primers targeting the left side of bait DSB and performed linear amplification to acquire prey sequences. Briefly, we used 5 μg sonicated DNA as input and amplified the target with high Phanta Max Super-Fidelity DNA Polymerase (Vazyme) using 5’-biotinylated primers, which can be captured efficiently by streptavidin beads and ease downstream enrichment. The linear amplification conditions are as follows: initial denaturation at 98°C for 3 min, 85 cycles of denaturation at 98°C for 30 sec, annealing at 58°C for 30 sec, and extension at 72°C for 90 sec, followed by a final extension at 72°C for 5 min. The linear amplification products were enriched with streptavidin beads. To get rid of free primers, we used BW buffer (5mM Tris-HCl, 0.5mM EDTA, 1M NaCl) to wash the beads. Finally, we resuspended the beads with ddH_2_O.

Considering various amplification 3’ ends, we ligated linear amplification products from the last step with annealed partial double-strand adaptors which have six random nucleotides at the 3’ end of one strand. After adaptor ligation, we proceeded with on-bead PCR using high Phanta Max Super-Fidelity DNA Polymerase (Vazyme) with P5/P7 adaptors. The PCR conditions were as follows: initial denaturation at 95°C for 5 min; 19 cycles of denaturation at 95°C for 30 sec, annealing at 60°C for 30 sec, extension at 72°C for 60 sec; followed by a final extension at 72°C for 5 min. The PCR products were purified with the High-Pure PCR Product Purification kit (Roche) and the library was sequenced by Illumina X ten platform.

### Customized computer program for reads processing

Although Cas9 has been reported to have staggered cleavage activity, up to now, there has not been any alignment software that takes this into account. We developed an alignment program that considers the complexity and diversity of Cas9 cleavage activity. With this program, we can obtain more precise alignments and thus ease downstream analyses.

CRIPSR-related insertions and deletions are frequently consecutive nucleotides. Software such as CrisprVariants (32) and AmpliconDIVider (33) maps next generation sequencing (NGS) reads by traditional aligners like BWA-MEM (34) and NovoAlign (http://www.novocraft.com). The software often reports CRISPR-unrelated short non-consecutive insertions and deletions. To solve this problem, Labun et al. developed ampliCan (35) by removing the gap-extension penalty and by modifying other scoring parameters of the Needleman-Wunsch algorithm. Thus, ampliCan tends to report consecutive long deletions and/or insertions. However, the indels reported by ampliCan are not promised to be at the Cas9 cleavage sites. Clement et al. proposed a partial solution to this problem in CRISPResso2 (36) by introducing a bonus at the cleavage site to incentivize indels there. Nevertheless, this does not completely solve the problem because the Cas9 cleavage may be staggered (20,24). In particular, it is not proper to treat the diverse profiles of Cas9 endonucleolytic cleavages as a single position of the -3 nucleotide upstream of PAM.

We develop a new program to solve this conundrum. It aligns each NGS input read to the junctional reference by two levels of optimization. Each NGS input read is separated into three parts before being mapped to the junctional reference. At the lower level, the program searches the optimal alignments of the left and right parts to the junctional reference, and the possibly-empty middle part is the unmapped random insertion. At the upper level, the program searches the optimal separation of the three parts. The two levels of optimization are technically integrated into dynamic programming. We permit the left and right parts of each NGS input read to overlap to capture the overhang of the staggered Cas9 cleavage ends. The detailed mathematical design and generalization as well as computational dynamic programming and source code (main.cpp) including its usage (Supplementary Notes S1-S4) are available on the GitHub/Gitee Platform (https://gitee.com/ljw20220910/Rearrangement).

### Calling for insertions and small deletions

Insertions and small deletions are called as previously described with optimizations (20,24). We designed PCR primer pairs near the junctional sites (Supplementary Table S1) for generating amplicon libraries (PCR products not more than 290bp in size) to assay small indels at junctions of chromosomal rearrangements. Therefore, paired end sequences can be merged for each read. In total, we obtained about 1.6 billion reads for this assay. After demultiplexing raw data of the FASTQ format with the index and barcode, we trimmed the sequences with Cutadapt (37). For each member of the amplicon library, we then merged the two paired-end sequence reads (read1 and read2) using PANDAseq (38). We divided each junctional repair outcome of chromosomal rearrangements into the four groups of deletions, insertions, indels, and precise ligations, and calculated their respective frequencies.

### Large resection analysis

Reads were mapped to the hg19 genome in both strands with our customized program. For reads covering large resections during DNA-fragment deletion, we required that the second segment maps strictly downstream of the first segment and that both are mapped to the forward strand. If the second segment maps to the reverse strand, then this is the case of large resections during DNA-fragment inversion.

### Mathematical estimation for MMEJ probability of small deletions

The region of the length *n* around a Cas9 cleavage has microhomology if and only if the length *M* of the longest common sequences flanking the cleavage site in the region is larger than a certain artificial threshold *L* defined by biological experiments. Although it is not easy to obtain the explicit cumulative probability distribution of *M*, an estimation is available (39) by transforming this microhomology problem within a region of DNA sequences into the problem of tossing a coin for a specific number of times equal to the length of DNA. However, each position of a DNA sequences can have any of the four bases of G,C,A,T in contrast to that each coin-tossing only has either head (obverse) or tail (reverse). The length *R* of the longest run of heads in the first *n* tosses of a coin is approximately log_2_(*n*) (40). More strictly, *R*/log_2_(*n*) converges to 1 almost everywhere as *n* tends to infinity.

By generalizing, Richard Arratia and Michael S. Waterman prove that *M*/log_1/*p*_(*n*) converges to 2 almost everywhere as *n* tends to infinity (39), where *p* = 1/4 is the probability that two random nucleotides in the corresponding position of the microhomology flanking the cleavage site are the same. For small *n*, they estimate the probability of *M* ≤ *L* with an upper bound 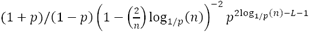and a lower bound (39).

We generated the curve of the lower bound estimations of MMEJ probabilities *P*(*M* ≥ 2) with increasing deletion sizes *n* for the panel of Figure 2I with a customized MATLAB script (Supplementary Note S5).

### Statistical analysis

All high-throughput sequencing libraries are constructed with at least two replicates. The significance tests are performed using the GraphPad software with Two-tailed t-tests, with one, two, three, and four asterisks indicating *P*-values less than 0.05, 0.01, 0.001 and 0.0001, respectively.

## RESULTS

### Reporter assay systems for large resections

In contrast to the small insertions from staggered Cas9 cleavages, there is little known about either small or large deletions. To provide mechanistic details into these processes, we first developed reporter assays using *HPRT1* (hypoxanthine phosphoribosyltransferase 1) and *DCK* (deoxycytidine kinase) systems. The *HPRT1* gene functions in the purine synthesis pathway and the encoded enzyme converts 6-thioguanine (6-TG) into a toxic product of thioguanine nucleotides. Thus, only *HPRT1*-defective cells can survive in 6-TG supplemented medium (Supplementary Figure S2A and B). The *DCK* gene encodes an essential enzyme for DNA synthesis. Therefore, *DCK*-defective cells cannot grow in normal medium (Supplementary Figure S2C and D). Thus, successful DSB events induced by Cas9 programmed with single sgRNAs targeting exons or with dual sgRNAs targeting introns of these genes can be readily assayed by cell growth in these two systems (Supplementary Figure S2).

More specifically, if we design single sgRNAs targeting exonic sequences (Supplementary Figure S2A and C), these two reporter systems can assay the efficiency of Cas9-induced DSB repair (Supplementary Figure S2B and D). If we design dual sgRNAs targeting exon-proximal intronic sequences (Supplementary Figure S2E and G), these reporter systems can be used to assay large resections into the flanking exonic sequences (Supplementary Figure S2F and H) because, with no large resection into the flanking exons, pre-mRNA splicing will not disrupt the normal function of *HPRT1* or *DCK*. We first examined the efficiency and sensitivity of these reporter systems (Supplementary Figure S2B, D, F, H) and indeed both the *HPRT1* and *DCK* reporter assay systems indicated that there exist large resections into the flanking exons for Cas9 programmed with dual sgRNAs (Supplementary Figure S2F and H).

### Polymerases in Cas9-induced large resections

We then used these reporter assay systems to investigate the roles of DNA polymerase genes (*PolM, PolD, PolL, PolQ*, and *PolK*) in Cas9-induced large resections. For the *HPRT1* reporter assay, upon *PolL* (encoding Polλ) or *PolQ* knockdown, especially *PolQ* knockdown, there is more cell growth despite adding 6-TG (Figure 1A and B), which suggested that Polθ and Polλ play a role in Cas9-induced large resections into the flanking exons of *HPRT1*. As a control, RT-PCR experiments demonstrated normal splicing in wild-type cells upon dual Cas9 cleavages within the intron 2 of *HPRT1* (Figure 1C). However, there are observable decreases of spliced *HPRT1* mRNA upon *PolL* or *PolQ* knockdown (Figure 1C), suggesting that there are increased large resections into the flanking *HPRT1* exons upon perturbation of *PolL* or *PolQ*. We made similar observations of increased large resections upon *PolL* or *PolQ* knockdown using the DCK reporter system (Figure 1D-F). DNA sequencing confirmed large resections from the second targeting site within intron 2 into the downstream exon 3 of *HPRT1* during DNA-fragment deletion (Figure 1G). In addition, we also confirmed large resections during DNA-fragment inversion (Figure 1H). Finally, there exist large resections at the upstream cleavage junction (Figure 1I). Together, these data suggest that both Polλ and Polθ play a role in Cas9-induced large resections programmed with dual sgRNAs.

**Figure 1.**
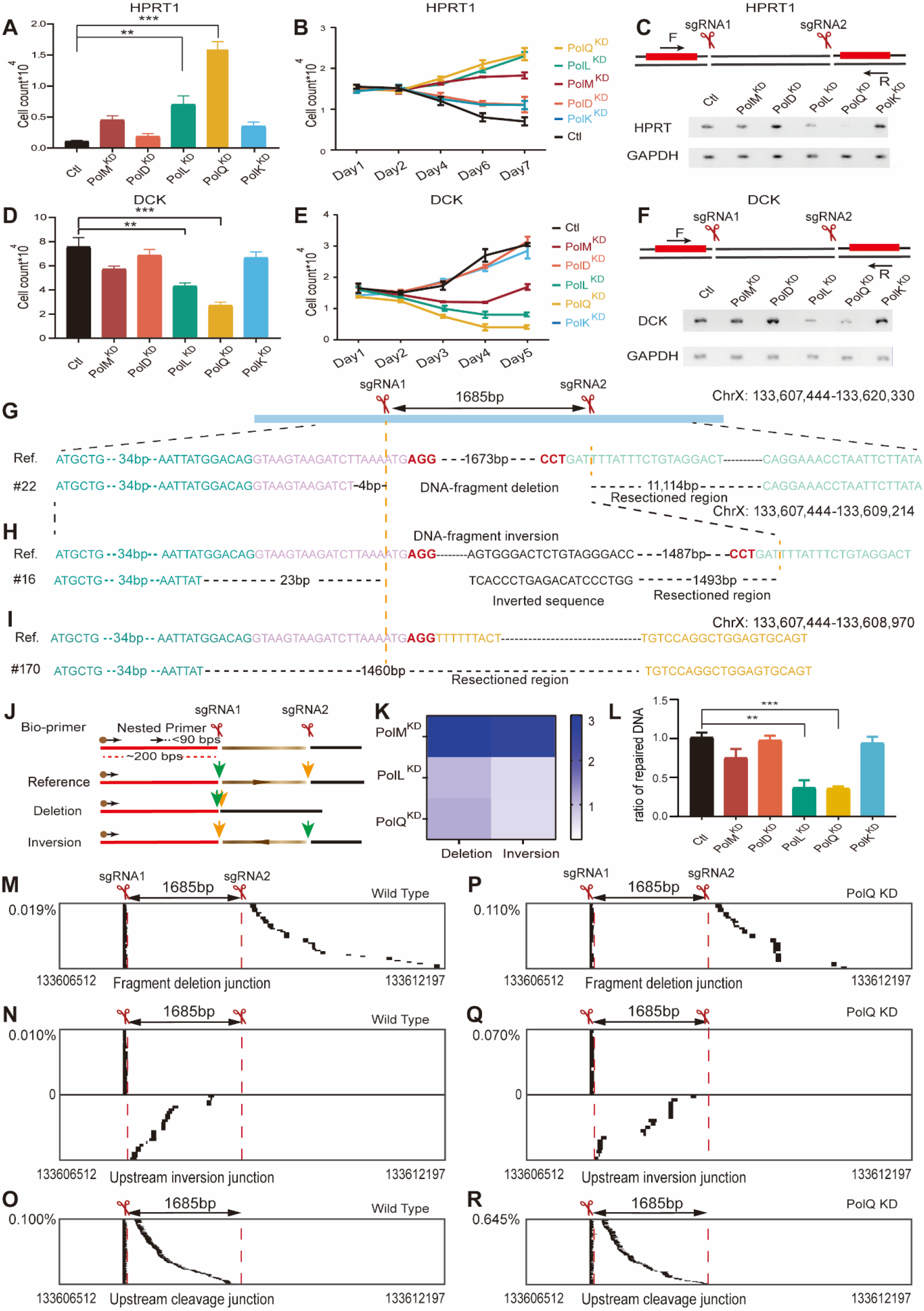
DNA polymerases in Cas9-induced large resections. Significant increases of resistance to 6-thioguanosine (**A, B**) and decreases of normal splicing (**C**) by Cas9-induced large resections of *HPRT1* upon knockdown of *PolQ* or *PolL*. Significant increases of sensitivity (**D, E**) and decreases of normal splicing (**F**) by Cas9-induced large resections of DCK upon knockdown of *PolQ* or *PolL*. Spliced *HPRT1* (**C**) and *DCK* (**D**) cDNAs are TA cloned and confirmed by Sanger sequencing in both orientations. Confirmation of large resections by DNA sequencing during DNA-fragment deletion (**G**) and inversion (**H**) as well as at the upstream cleavage junction (**I**) programmed by Cas9 with dual sgRNAs. Schematic (**J**) of LAM-HTGTS with biotinylated and nested primers and simultaneous assessment of DNA-fragment deletion and inversion (**K**) by next-generation sequencing (NGS). Significant decreases of repaired DNA upon knockdown of *PolQ* or *PolL* (**L**). Significant increases of Cas9-induced large resections during DNA-fragment deletion (**M, P**) and inversion (**N, Q**) as well as at the upstream cleavage junction (**O, R**) assayed by NGS upon knockdown of *PolQ* (**P**-**R**).

We then adopted LAM-HTGTS (31) to assay large resections during chromosomal rearrangements induced by Cas9 with dual sgRNAs. LAM-HTGTS can assay both DNA-fragment deletions and inversions simultaneously (Figure 1J). We observed a higher frequency of DNA-fragment deletions compared with inversions during DNA-fragment editing (Figure 1K). We also found that knockdown of *PolQ* or *PolL* results in significant decreases in repaired DNAs, suggesting again that Polθ and Polλ are required for DSB repairs during DNA-fragment editing (Figure 1L).

Computational analyses of the next-generation sequencing (NGS) data with a customized computer program (see reads processing in MATERIALS AND METHODS and Supplementary Notes 1-3) identified rare but significant portion of high throughput sequencing reads for the large resections during DNA-fragment deletions (Figure 1M). We also identified a large number of reads for the large resections that occurred during DNA-fragment inversions (Figure 1N). Finally, there exist a large number of reads of large resections at the upstream cleavage junctions (Figure 1O). These data demonstrated that there are asymmetrical large resections at the Cas9 cleavage site. Importantly, *PolQ* knockdown results in significant increases of large resections at all of these chromosomal rearrangement junctions (Figure 1P-R), in line with observed increases of large resections by the *HPRT1* and *DCK* reporter assay systems (Figure 1A-F). We also observed that the vast majorities of sequencing reads at junctions of chromosomal rearrangements have small indels, and that *PolQ* knockdown exhibits a more pronounced effect on chromosomal rearrangements programmed with dual sgRNAs (Supplementary Figure S3A, B, D, E) than on editing outcomes from single cleavages (Supplementary Figure S3C and F).

### Polθ in small deletions

In addition to rare large resections, small deletions are more frequently observed at junctions of chromosomal rearrangements during DNA-fragment editing. We found that upon *PolQ* but not *PolD* or *PolK* knockdown, there are consistent and significant decreases of small deletions at both upstream and downstream junctions of fragment inversions (Figure 2A and B) as well as at the junctions of fragment deletions (Figure 2C) and duplications (Figure 2D) at the *MeCP2* locus. We then examined the role of *PolQ* at four additional loci (namely, *MAZ, PRDM5, PARP1*, and *YY1*). *PolQ* knockdown results in significant decreases of small deletions at junctions of DNA-fragment editing in all of these four loci (Supplementary Figure S4). Taken together, these data suggest that Polθ is essential for generating small deletions during DNA-fragment editing by Cas9 with dual sgRNAs.

**Figure 2.**
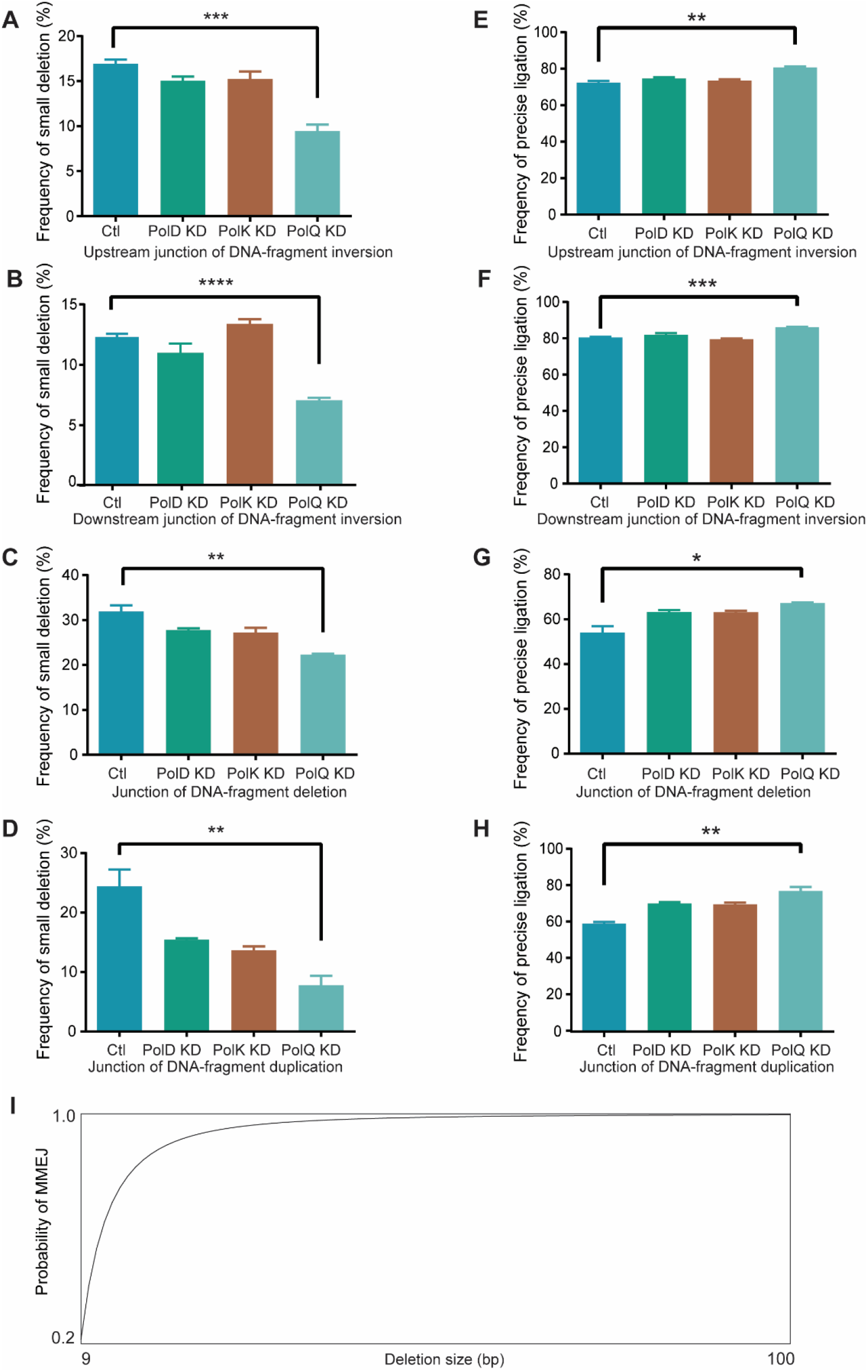
Polθ promotes small deletions of chromosomal rearrangements. Significant decreases of small deletions and increases of precise ligations at the upstream (**A, E**) and downstream (**B, F**) junctions of DNA-fragment inversions as well as at the junctions of DNA-fragment deletions (**C, G**) and duplications (**D, H**) upon knockdown of PolQ. (**I**) Estimation of the probability of MMEJ with increasing deletion size.

Disruption of *CtIP* or *FANCD2*, two DNA repair genes required in the Alt-NHEJ pathway, results in increased precise ligations at junctions of chromosomal rearrangements during DNA-fragment editing (24). Thus, cNHEJ, which functions in precise ligations of this editing, competes with Alt-NHEJ for repair substrates. Interestingly, *PolQ* knockdown results in a consistent increase of precise ligations at both upstream and downstream junctions of inversions (Figure 2E and F) as well as junctions of deletions (Figure 2G) and duplications (Figure 2H) at the *MeCP2* locus. In addition, *PolQ* knockdown also results in increased precise ligations in the *MAZ, PRDM5*, and *PARP1* loci (Supplementary Figure S5). This is in line with the competition of Alt-NHEJ and cNHEJ for repairing Cas9-induced DSB ends during DNA-fragment editing.

Small deletions are editing outcomes of the MMEJ repair pathway upon Cas9 cleavages. The size of MMEJ deletions is determined by the distance from embedded microhomology to the Cas9 cleavage site. Computational analysis revealed that the conservative estimation of the probability of finding at least 2bp microhomology increases rapidly to 99.7% as the deletion size reaches 100bp (Figure 2I). Accordingly, we analyzed small deletions of less than 100bp in detail below.

### Distinct mechanisms for CRISPR 1bp and >1bp deletions

Recent gene editing using Cas9-Pol I fusion proteins revealed that 1bp and >1bp deletions may be generated differently (41), but the underlying mechanism is unknown. To this end, we separately analyzed 1bp and 2-100bp deletions. Remarkably, upon *PolQ* knockdown, there is a significant increase in 1bp deletions during DNA-fragment editing at the *MAZ* locus (Figure 3A). In contrast, *PolQ* knockdown results in a significant decrease of 2-100bp deletions at the *MAZ* locus (Figure 3B and C). We performed *PolQ* knockdown experiments for four additional loci (namely, *MeCP2, PARP1, PRDM5*, and *YY1*), and observed similar increases of 1bp deletions and decreases of 2-100bp deletions (Figure 3D-O). These data suggest that Polθ is essential for the generation of small deletions of 2-100bp, which most likely resulted from the processing by the MMEJ pathways during CRISPR DNA-fragment editing. By contrast, the mechanism for generating 1bp deletions is different, most likely resulting from the processing by the cNHEJ pathway during the DNA-fragment editing.

**Figure 3.**
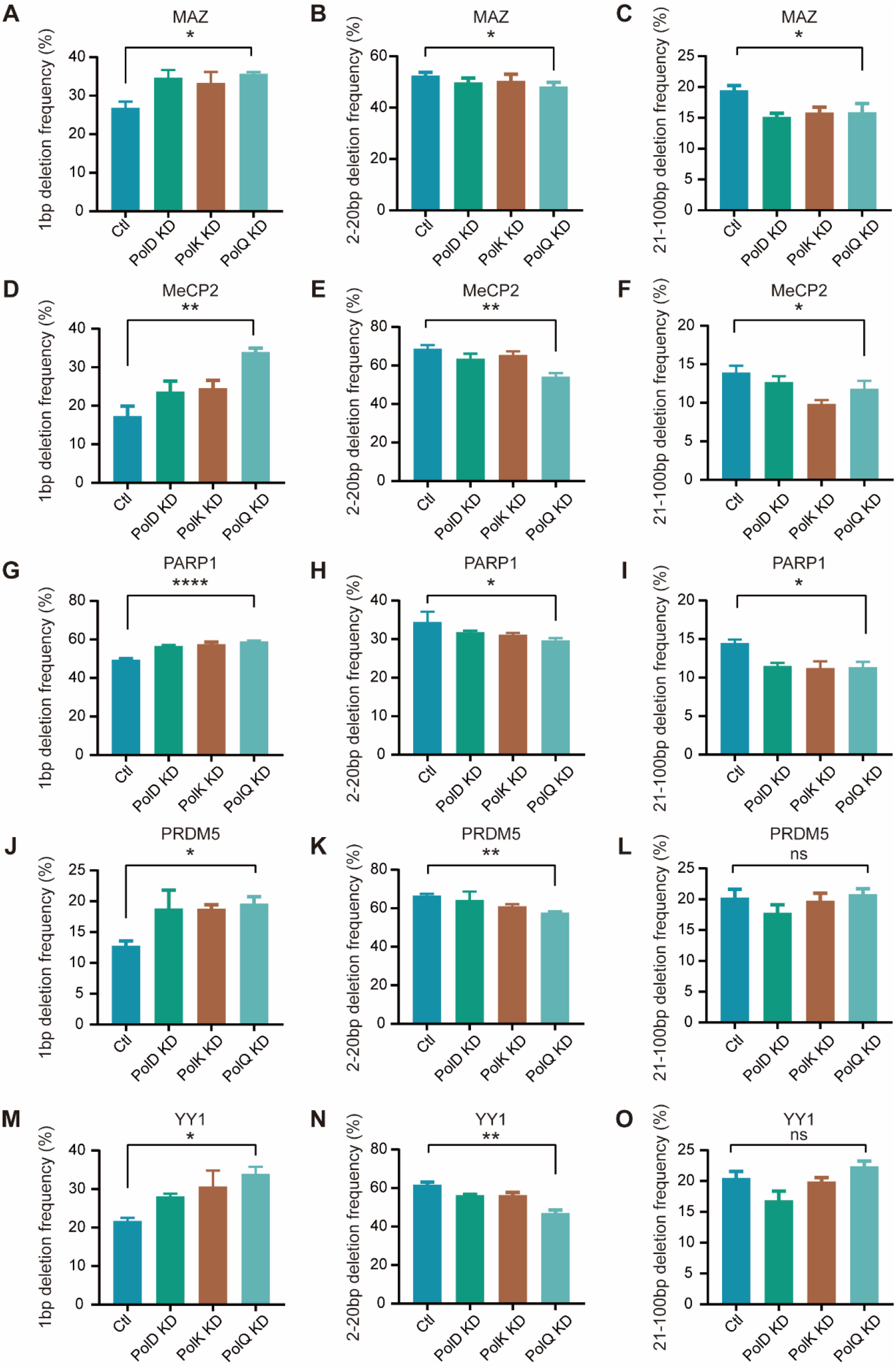
Mechanistic differences for 1bp and 2-100bp deletions. Significant increases of the frequency of 1bp deletions (**A, D, G, J, M**) and decreases of the frequency of 2-100bp deletions (**B, E, H, K, N** and **C, F, I, L, O**) upon *PolQ* knockdown at the *MAZ* (**A**-**C**), *MeCP2* (**D**-**F**), *PARP1* (**G**-**I**), *PRDM5* (**J**-**L**), and *YY1* (**M**-**O**) loci.

### Polλ in 1bp deletions

To provide further insight into the molecular mechanisms underlying 1bp and >1bp deletions, we knocked down two members of the DNA polymerase X family, *PolL* and *PolM*, separately or together. Interestingly, we found that *PolL* knockdown results in a significant decrease of 1bp deletion frequency (Supplementary Figure S6), suggesting an essential role of Polλ in the 1bp deletions. In contrast, *PolL* knockdown leads to a significant increase of >1bp deletion frequency (Supplementary Figure S6). This again suggests that 1bp and >1bp deletions are generated by the different repair pathways of cNHEJ and MMEJ, respectively. Further knockdown of *PolM*, another member of the polymerase X family, in combination with the *PolL* knockdown only produced minimal effects (Supplementary Figure S6). This suggests that Polμ, in contrast to Polλ, plays a limited role in the generation of small deletions during CRISPR DNA-fragment editing. In conjunction with the data from the *PolQ* knockdown (Figure 3), we conclude that the mechanism of generating 1bp deletions is fundamentally different from that of generating >1bp deletions, and that they are generated by cNHEJ and MMEJ pathways, respectively.

### Polλ fill-in of Cas9 staggered cleavages

Recent studies have revealed that Cas9 cleavage generates staggered DSB ends with 1-3bp 5’ overhangs in addition to blunt ends during chromosomal rearrangements induced by Cas9 with dual sgRNAs (1,20,24). Consistent with staggered endonucleolytic Cas9 cleavage, we found that 1-3bp deletions at junctions of chromosomal rearrangements are strongly biased toward the -4, -5, and -6 positions upstream of the PAM site (Supplementary Figure S7). However, the mammalian polymerase(s) responsible for the fill-in of the staggered Cas9 DSB ends is presently unclear.

We thus analyzed 1-3bp templated insertions from fill-in of staggered DSB ends upon polymerase knockdown. However, the available software to characterize Cas9 editing does not take its staggered cleavage into account (32,33,35,36). To this end, we developed a customized computer program to specifically enable this analysis (see reads processing in MATERIALS AND METHODS and Supplementary Notes 1-3).

We first analyzed the 1-3bp templated insertions from staggered Cas9 cleavages with sgRNA1 and sgRNA2 at the *MAZ* locus and found that templated 1-3bp insertions generated by both sgRNA1 and sgRNA2 are significantly decreased upon knockdown of *PolL*, and to a lesser extent upon knockdown of *PolM* (Figure 4A and B). However, knockdown of both *PolL* and *PolM* together does not lead to further decreases in templated 1-3bp insertions compared to the knockdown of *PolL* only (Figure 4A and B). This suggests that Polλ has a dominant role in the fill-in of staggered Cas9 cleavages, consistent with its role in promoting mutagenesis observed in a recent CRISPR large-scale analysis (42). We then performed these knockdown experiments for four more loci (*PARP1, PRDM5, YY1*, and *MeCP2*) and found significant decreases of templated 1-3bp insertions for *PARP1, PRDM5*, and *YY1* upon knockdown of *PolL* (Figure 4C-H). The data for the *MeCP2* locus is not significant as the cleavage profile of *MeCP2* is strongly biased toward blunt cleavages owning to the predominant precise ligations at junctions of chromosomal rearrangements in the *MeCP2* locus (Figure 2). Finally, we found no significant difference in frequency of 1-3bp templated insertions upon knockdown of *PolD, PolK*, or *PolQ* (Supplementary Figure S8). Altogether, we conclude that Polλ is the main polymerase responsible for the cellular fill-in of staggered Cas9 endonucleolytic cleavages *in vivo*.

**Figure 4.**
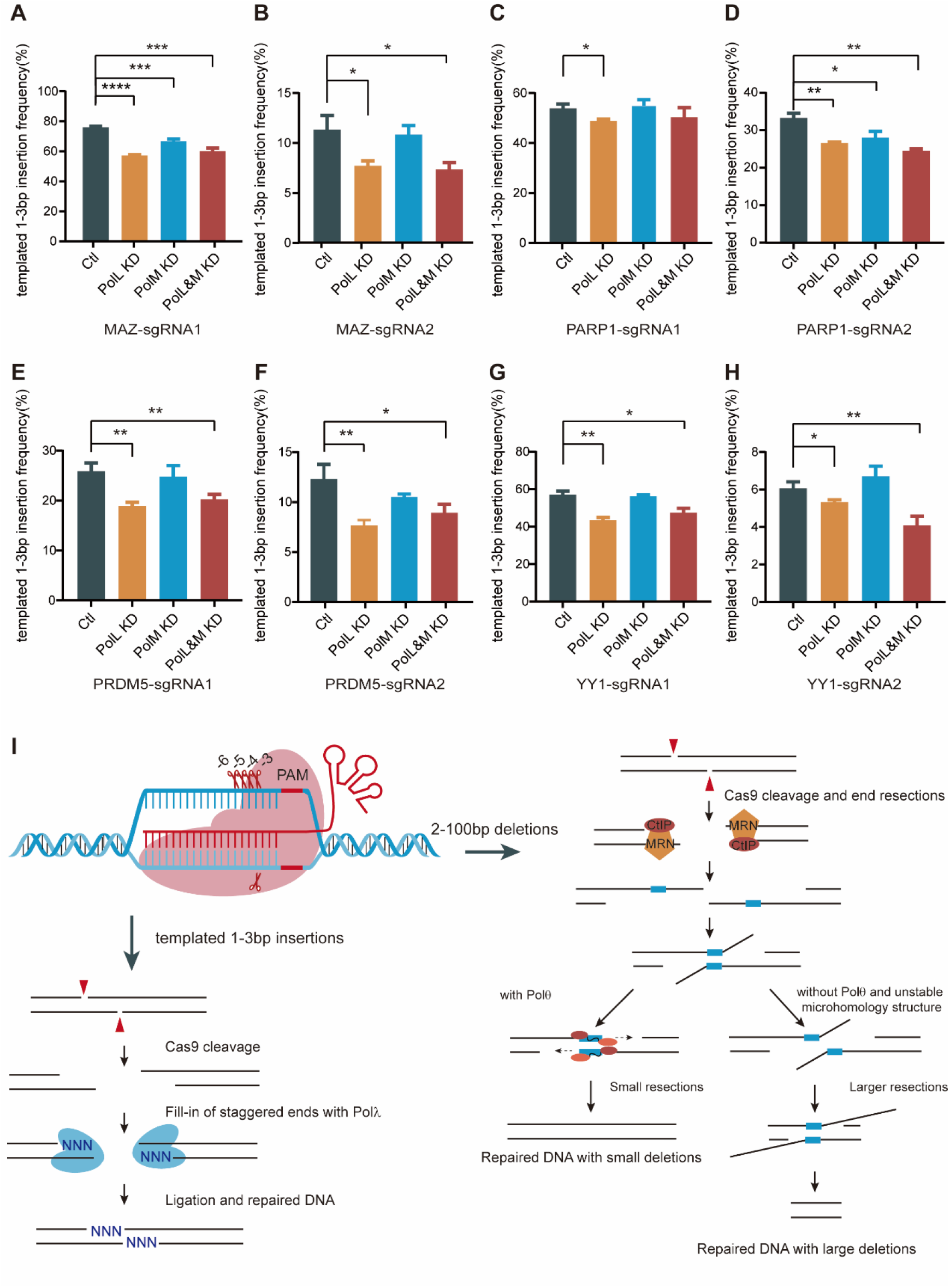
Fill-in of staggered Cas9 DSB ends by Polλ. Significant decreases of templated 1-3bp insertions at Cas9 cleavage junctions programmed with dual sgRNAs at the *MAZ* (**A, B**), *PARP1* (**C, D**), *PRDM5* (**E, F**), and *YY1* (**G, H**) loci upon *PolL* knockdown. Schematic of the role of Polλ and Polθ in repairing Cas9-induced DSB ends (**I**).

## DISCUSSION

DNA polymerases are thought to counteract nuclease activities of the MRN complex during CRISPR/Cas9 gene editing (41). Here we have identified for the first time the polymerases involved in CRISPR gene editing events, providing important mechanistic details about various deletions and insertions during DNA-fragment editing. Overall, we find that Polλ, and to a lesser extend Polμ, are the long-sought DNA polymerases that fill-in the staggered DSB ends from programmed Cas9 cleavage. Surprisingly, we find prominent large resections at junctions of chromosomal rearrangements. In addition, we find that Polλ and Polθ are very important for these large resections. We also find that mechanisms for 1bp and >1bp deletions are distinct because of opposite dependencies on Polλ and Polθ in generating small deletions. Hence, there appears to be fundamentally different pathways enlisted in these Cas9-dependent genome modifications.

Although we find a role of Polλ in large resections induced by Cas9 with dual sgRNAs (Figure 1A-F), the underlying mechanism is still not clear. In particular, since Polλ is a member of family X polymerases, and it participates in cNHEJ via BRCT interaction with the Ku-XRCC4-LIG IV complex (5), it may be able to block large resections. However, how Polλ precisely blocks large resection remains to be investigated in the future.

We find prominent numbers of sequencing reads indicating complex large resections at junctions of chromosomal rearrangements during CRISPR DNA-fragment editing by Cas9 programmed with dual sgRNAs. Previous studies revealed that Polθ can mediate the joining of two 3’ overhangs with 2-20bp microhomology exposed after MRN resection (43). Specifically, Polθ facilitates microhomology search and stabilizes annealing of microhomologous sequences via its complex activities such as dNTP-dependent 3’-end trimming and template-dependent DNA synthesis (44). Here, we find that large resections are increased and small deletions are decreased upon *PolQ* knockdown, which suggests the essential role of Polθ in suppressing Cas9-mediated large resections and in inducing MMEJ with small deletions. It is possible that Polθ perturbation impairs MMEJ which may permit continuous resection into flanking regions thus resulting in large resections.

It is interesting that knockdown of *PolQ* and *PolL* reveals mechanistic differences of generating 1bp and >1bp deletions during DNA-fragment editing. It is consistent with that NHEJ is responsible for generating 1b deletion and MMEJ using microhomology sequences embedded in the flanking region results small deletions of 2-100bp. However, the exact locations of microhomology are sequence context dependent. In addition, the NHEJ repair pathway may generate deletions of 1bp or very few base pairs and deletions of 2bp or 3bp may not be generated by the MMEJ pathway. The exact turning point between NHEJ and MMEJ may be locus dependent. Finally, we find that Polλ is the main cellular polymerase responsible for the fill-in of staggered Cas9 cleavages *in vivo*. Taken together, our data reveal the crucial role of Polλ and Polθ in the repair of Cas9-induced DSBs (Figure 4I) and should be conducive to the development of controllable CRISPR chromosomal rearrangements.

## Supporting information

All supplementary information

## DATA AVAILABILITY

High-throughput sequencing files have been submitted into to SRA with accession number SRP405576.

## ACKNOWLEDGEMENT

We thank Prof. Dan Czajkowsky for great improvements on the manuscript and all members of our laboratory for helpful discussion.

