## Supplementary material for "DNA Polymerases in Precise and Predictable CRISPR/Cas9-mediated Chromosomal Rearrangements": All supplementary information

### This file includes eight figures, one table, and five notes:

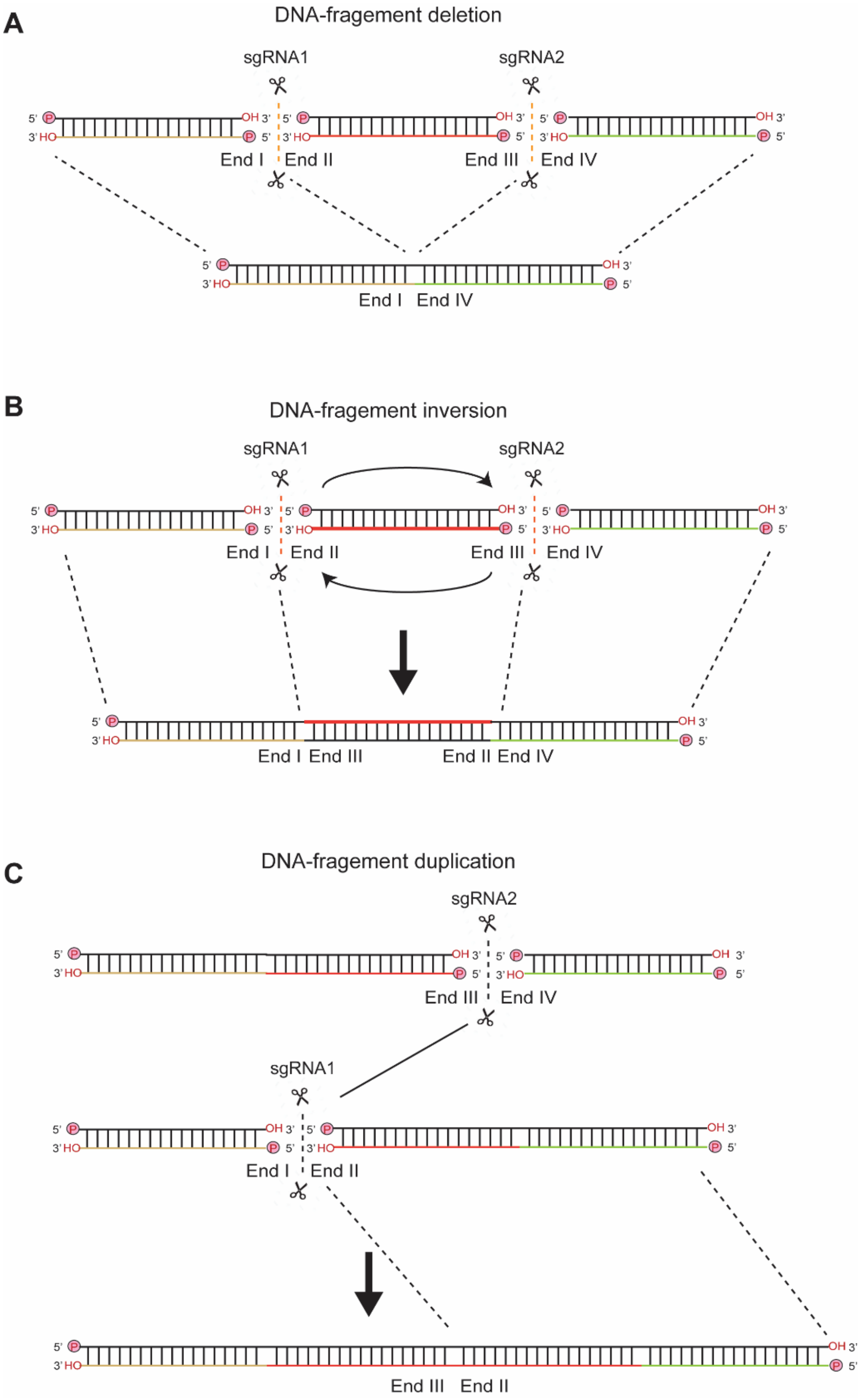


**Supplementary Figure S1.** DNA-fragment editing by Cas9 with dual sgRNAs results in chromosomal rearrangements including DNA-fragment deletion, inversion, and duplication. Cleavages of Cas9 programed with dual sgRNAs result in two cuts with four double-stranded break (DSB) ends: I, II, III, and IV. (**A**) Ligation of DSB ends I and IV leads to DNA-fragment deletion. (**B**) Ligation of DSB ends I and III as well as ends II and IV leads to DNA-fragment inversion. (**C**) Transallelic ligation of DSB ends III with II leads to DNA-fragment duplication.

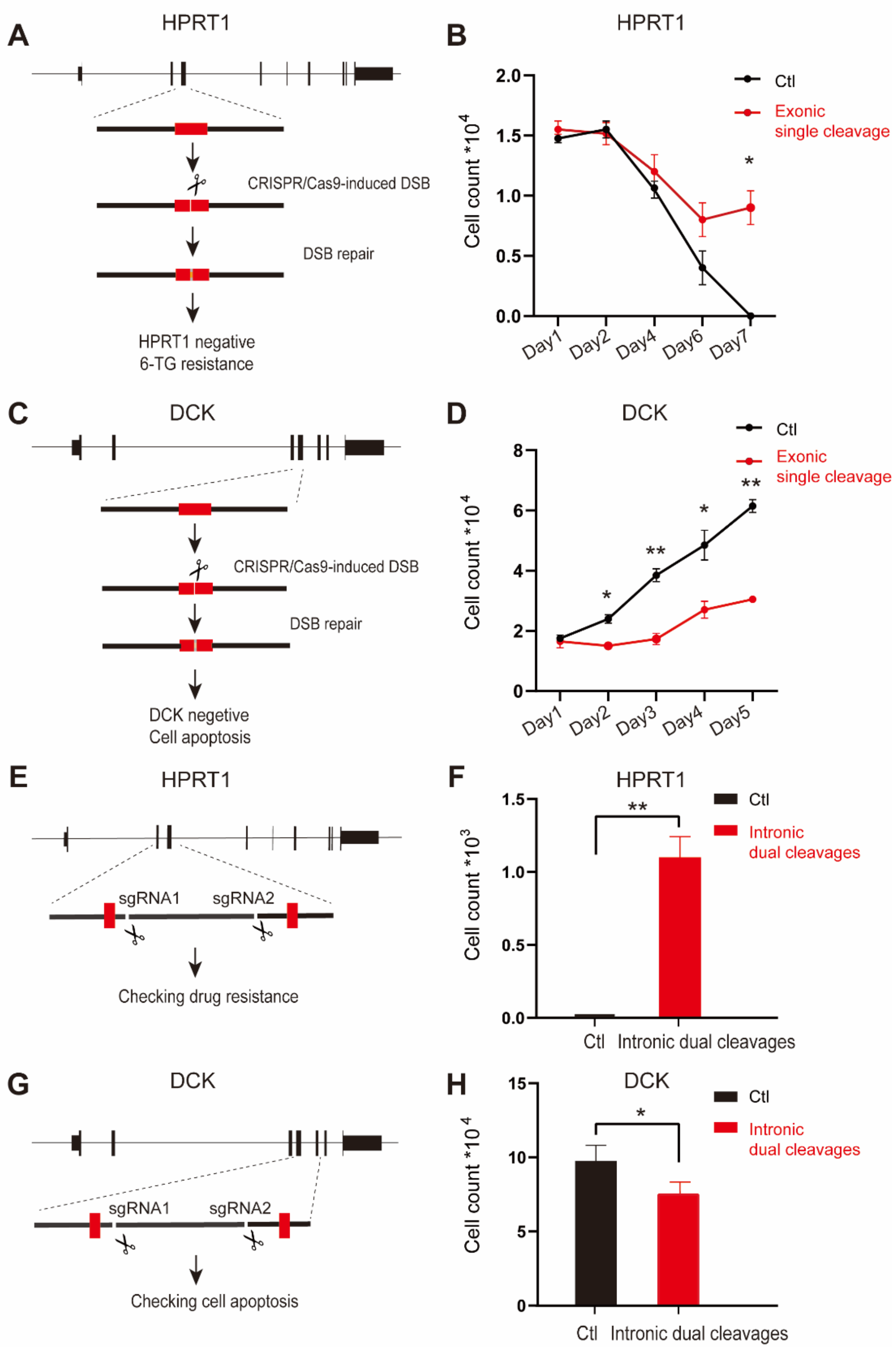


**Supplementary Figure S2.** *HPRT1* and *DCK* reporter assay systems for Cas9-induced large resections. Schematics (**A**, **C**) and cell growth (**B**, **D**) of Cas9 with single sgRNAs targeting exonic sequences of *HPRT1* (with 6-TG) and *DCK*. Schematics (**E**, **G**) and cell growth (**F**, **H**) of Cas9 with dual sgRNAs targeting exon-proximal intronic sequences of *HPRT1* (with 6-TG) and *DCK*. Normal splicing of the intron with no large resection will not perturb *HPRT1* or *DCK* gene expression. Cas9-induced large resections into the flanking exonic sequences will disrupt normal splicing and perturb *HPRT1* or *DCK* gene function.

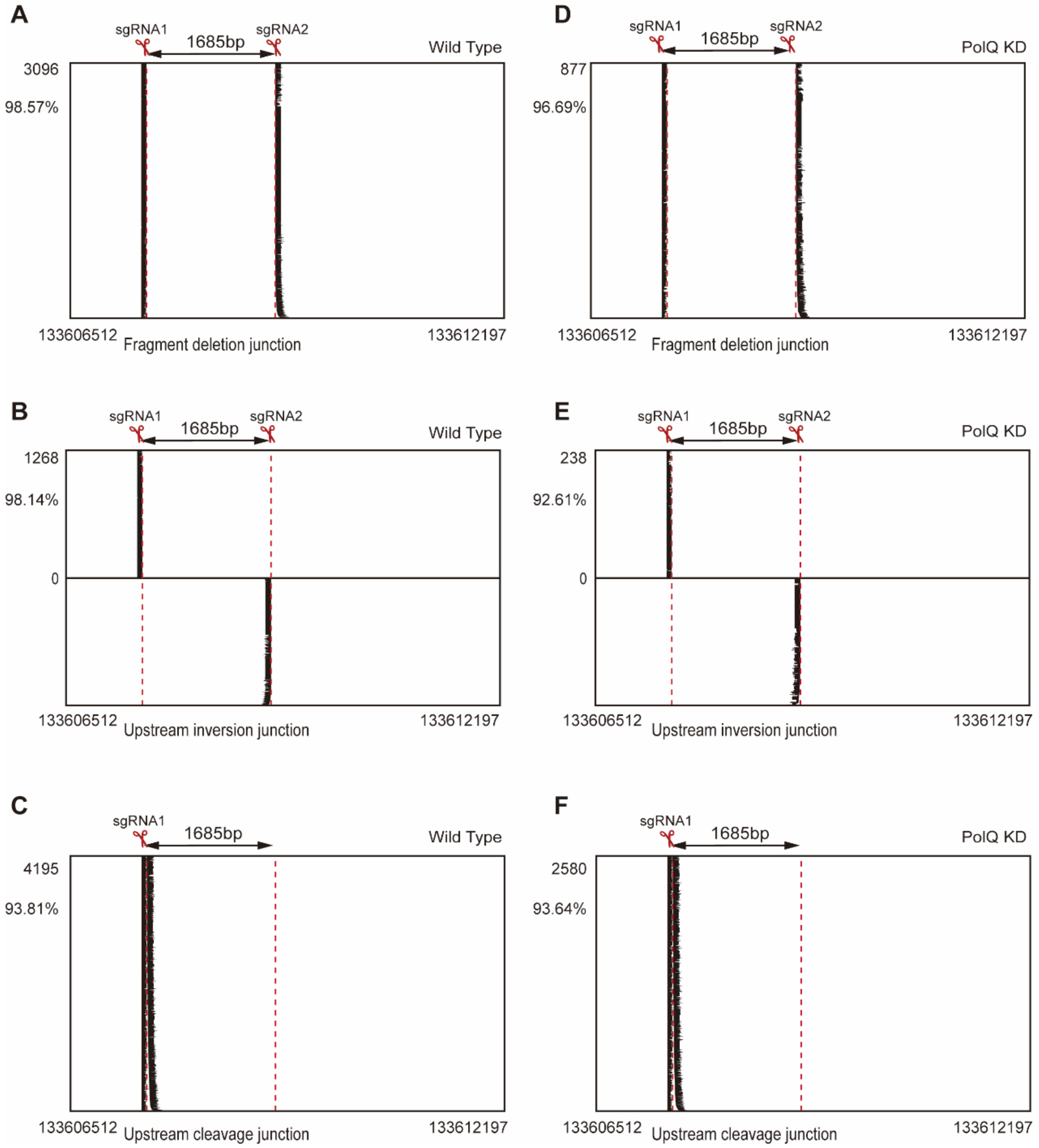


**Supplementary Figure S3.** High-throughput NGS of junctional sequences of DNA-fragment deletions (**A**, **D**) and inversions (**B**, **E**) as well as of single cleavage junctions of Cas9 with sgRNA1 (**C**, **F**) upon *PolQ* knockdown. Note the more pronounced effects on chromosomal rearrangement junctions of DNA-fragment deletions and inversions (**A**, **B**, **D**, **E**) than on single cleavage junctions (**C**, **F**).

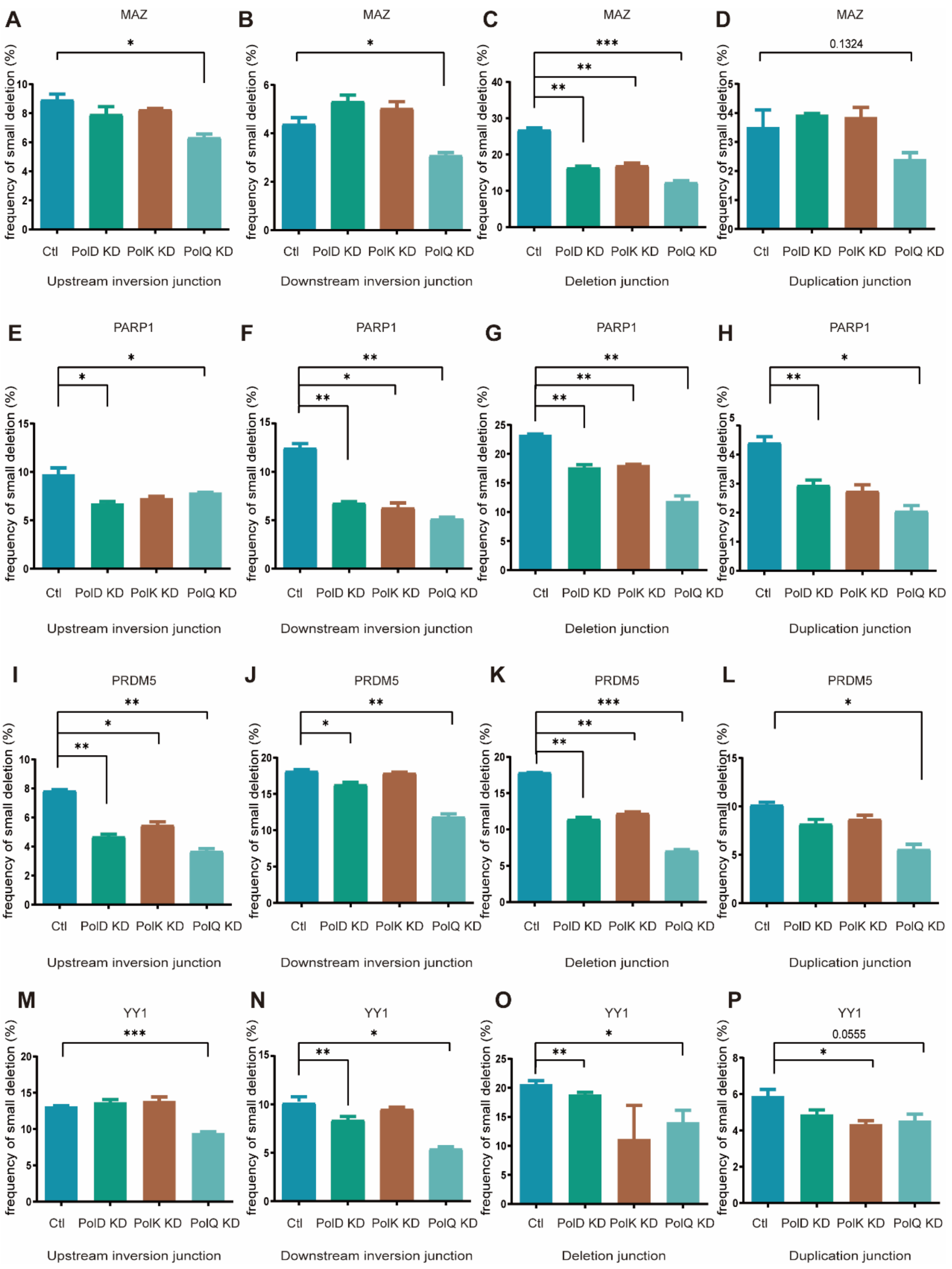


**Supplementary Figure S4.** Significant decreases in the frequency of small deletions at upstream (**A**, **E**, **I**, **M**) and downstream (**B**, **F**, **J**, **N**) junctions of DNA-fragment inversion as well as at junctions of DNA-fragment deletion (**C**, **G**, **K**, **O**) and duplication (**D**, **H**, **L**, **P**) at the *MAZ* (**A-D**), *PARP1* (**E-H**), *PRDM5* (**I-L**), and *YY1* (**M-P**) loci upon *PolQ* knockdown.

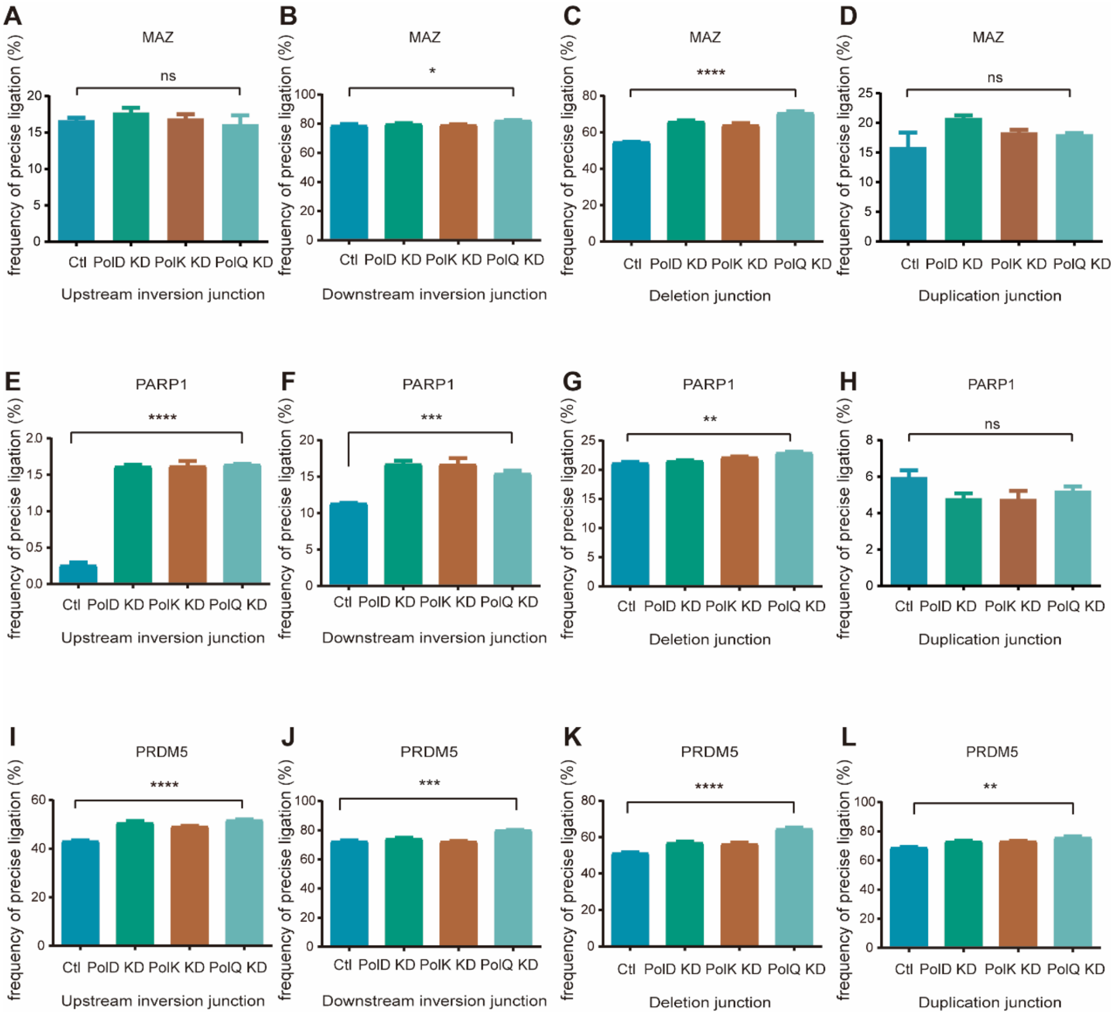


**Supplementary Figure S5.** Significant increases in the frequency of precise ligations at upstream (**A**, **E**, **I**) and downstream (**B**, **F**, **J**) junctions of DNA-fragment inversion as well as at junctions of DNA-fragment deletion (**C**, **G**, **K**) and duplication (**D**, **H**, **L**) at the *MAZ* (**A**-**D**), *PARP1* (**E**-**H**), and *PRDM5* (**I**-**L**) loci upon *PolQ* knockdown.

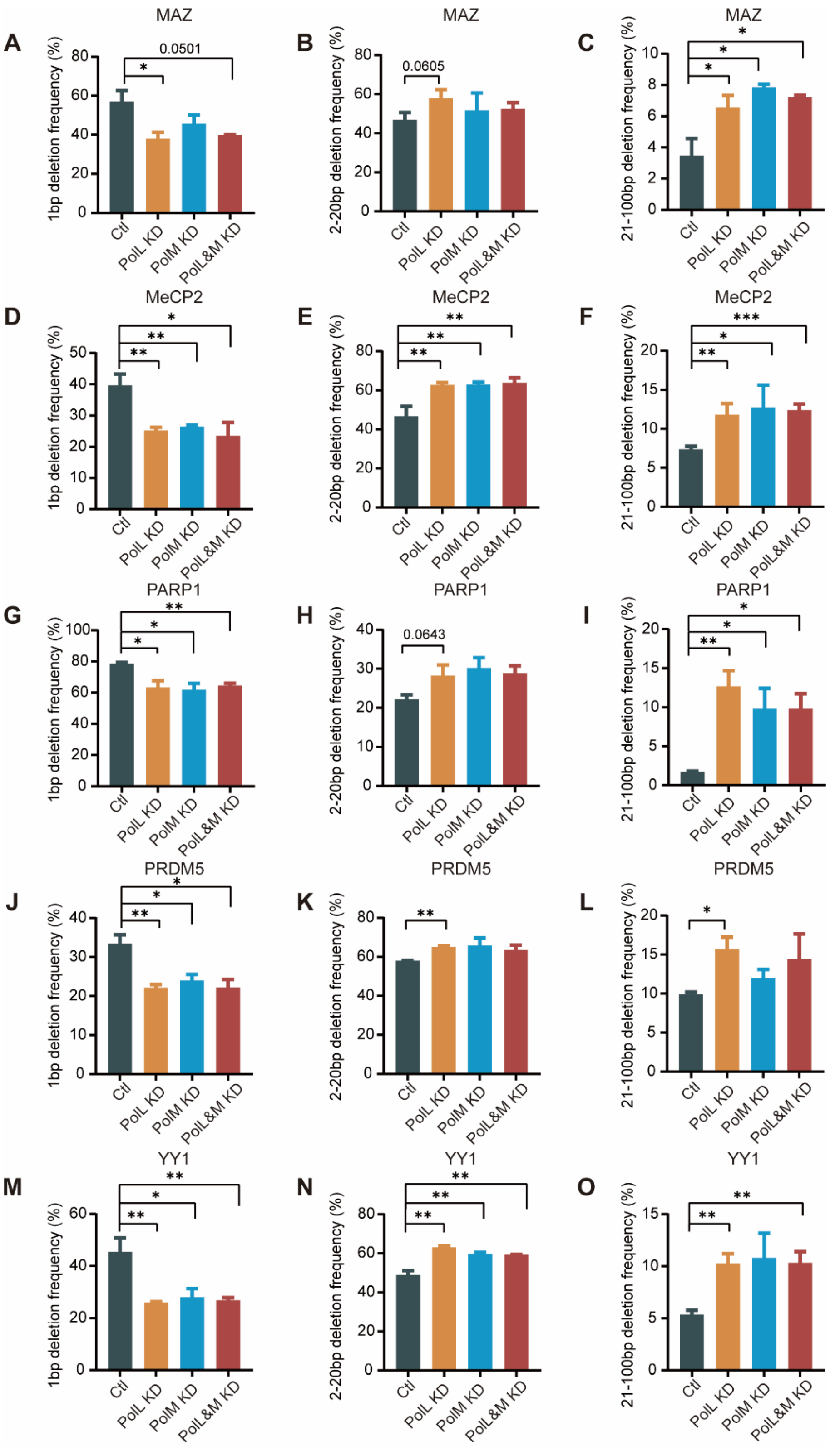


**Supplementary Figure S6.** Polλ enhances editing outcomes of 1bp deletions and suppresses the generation of >1bp deletions. There is a significant decrease of frequencies of 1bp deletions but an increase of frequencies of 2-20bp and 21-100bp deletions upon *PolL* knockdown at the *MAZ* (**A**-**C**), *MeCP2* (**D**-**F**), *PARP1* (**G**-**I**), *PRDM5* (**J**-**L**), and *YY1* (**M**-**O**) loci.

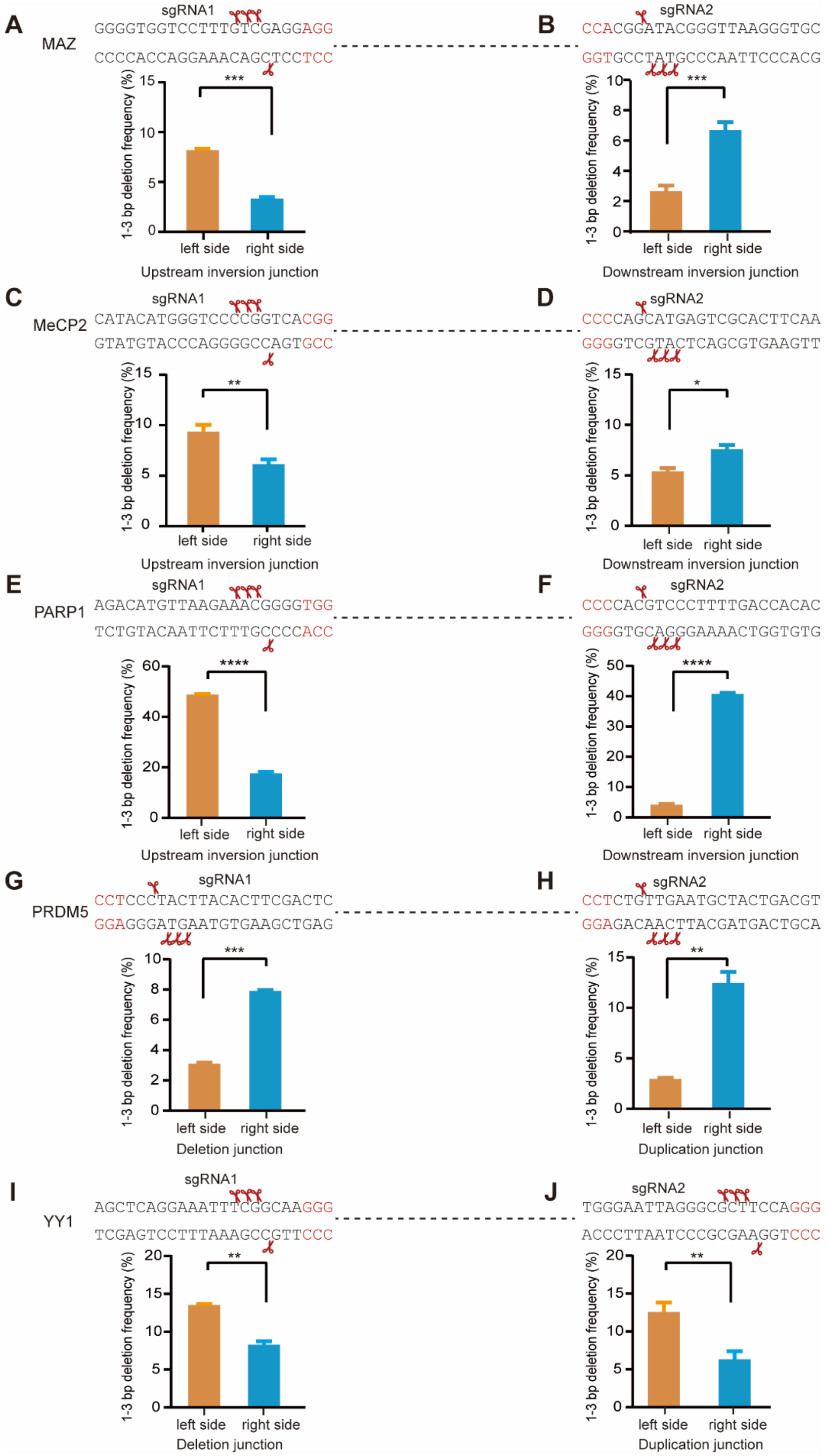


**Supplementary Figure S7.** Biased deletion of nucleotides at junctional sites of chromosomal rearrangements confirms the staggered Cas9 cleavages. 1-3bp deletion frequencies at the chromosomal rearrangement junctions are always biased toward -4, -5, and -6 positions upstream of the PAM site of both sgRNAs at the *MAZ* (**A**, **B**), *MeCP2* (**C**, **D**), *PARP1* (**E**, **F**), *PRDM5* (**G**, **H**) and *YY1* (**I**, **J**) loci.

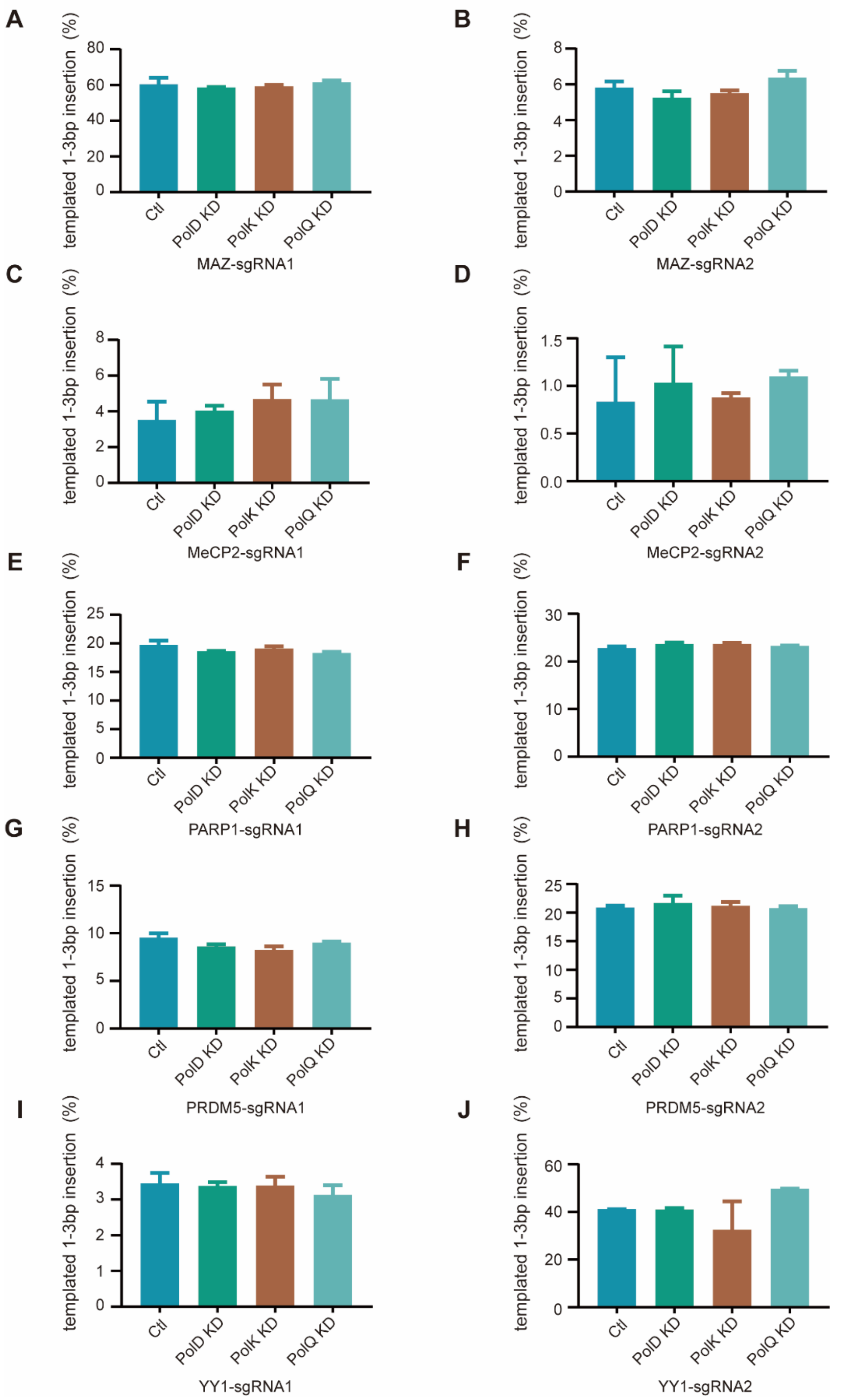


**Supplementary Figure S8.** Polδ, Polκ, and Polθ are not engaged in the fill-in of staggered Cas9 cleavage ends. There are no significant differences of 1-3bp templated insertions at the junctional sites of chromosomal rearrangements programmed with Cas9 and dual sgRNAs at the *MAZ* (**A**, **B**), *MeCP2* (**C**, **D**), *PARP1* (**E**, **F**), *PRDM5* (**G**, **H**), and *YY1* (**I**, **J**) loci upon knockdown of *PolD*, *PolK*, and *PolQ*.

**Supplementary Table S1.** Oligonucleotide sequences used in this study.

| | |
|---|---|
| Oligos for knocking down polymerases | |
| PL-E5-F | ACCGAGGCTCACAATCTCCCTCAA |
| PL-E5-R | AAACTTGAGGGAGATTGTGAGCCT |
| PL-E7-F | ACCGCTTCCTGGAACGTATGCCCA |
| PL-E7-R | AAACTGGGCATACGTTCCAGGAAG |
| PM-E10-F | ACCGGCTGCGCCGCTTCAGCCGGA |
| PM-E10-R | AAACTCCGGCTGAAGCGGCGCAGC |
| PM-E7-F | ACCGTACATCGGACCGCAGGACTG |
| PM-E7-R | AAACCAGTCCTGCGGTCCGATGTA |
| PQ-E16-F | ACCGACGCTCCAGAGTCTTTCAGG |
| PQ-E16-R | AAACCCTGAAAGACTCTGGAGCGT |
| PQ-E15-F | ACCGTTAGTAAGGATACCCGAACC |
| PQ-E15-R | AAACGGTTCGGGTATCCTTACTAA |
| PD1-E20-1F | ACCGGTATGGGAAGTAGACCTGGG |
| PD1-E20-1R | AAACCCCAGGTCTACTTCCCATAC |
| PD1-E22-1F | ACCGTGATGATCACGTAGGGGACG |
| PD1-E22-1R | AAACCGTCCCCTACGTGATCATCA |
| RT-PCR primers for the detection of large resections | |
| HPRT1-F | TGCTGACCTGCTGGATTACA |
| HPRT1-R | ACAATCCGCCCAAAGGGAAC |
| DCK-F | AGAGAAGCTGCCCGTCTTTC |
| DCK-R | CAGGAGCCAGCTTTCATGTT |
| GAPDH-F | GGAGTCCACTGGCGTCTTCAC |
| GAPDH-R | GCAGGAGGCATTGCTGATGAT |
| sgRNA sequences for the induction of chromosomal rearrangements | |
| MeCP2-sgRNA1-F | ACCGCATACATGGGTCCCCGGTCA |
| MeCP2-sgRNA1-R | AAACTGACCGGGGACCCATGTATG |
| MeCp2-sgRNA2-F | ACCGTTGAAGTGCGACTCATGCTG |
| MeCP2-sgRNA2-R | AAACCAGCATGAGTCGCACTTCAA |
| MAZ-sgRNA1-F | ACCGGGGGTGGTCCTTTGTCGAGG |
| MAZ-sgRNA1-R | AAACGGGGTGGTCCTTTGTCGAGG |
| MAZ-sgRNA2-F | ACCGGCACCCTTAACCCGTATCCG |
| MAZ-sgRNA2-R | AAACCGGATACGGGTTAAGGGTGC |
| PARP1-sgRNA1-F | ACCGAGACATGTTAAGAAACGGGG |
| PARP1-sgRNA1-R | AAACCCCCGTTTCTTAACATGTCT |
| PARP1-sgRNA2-F | ACCGGTGTGGTCAAAAGGGACGTG |
| PARP1-sgRNA2-R | AAACCACGTCCCTTTTGACCACAC |
| PRDM5-sgRNA1-F | ACCGGAGTCGAAGTGTAAGTAGGG |
| PRDM5-sgRNA1-R | AAACCCCTACTTACACTTCGACTC |
| PRDM5-sgRNA2-F | ACCGACGTCAGTAGCATTCAACAG |
| PRDM5-sgRNA2-R | AAACCTGTTGAATGCTACTGACGT |

| | |
|---|---|
| YY1-sgRNA1-F | ACCGAGCTCAGGAAATTTCGGCAA |
| YY1-sgRNA1-R | AAACTTGCCGAAATTTCCTGAGCT |
| YY1-sgRNA2-F | ACCGTGGGAATTAGGGCGCTTCCA |
| YY1-sgRNA2-R | AAACTGGAAGCGCCCTAATTCCCA |
| Oligos for the construction libraries for high throughput sequencing | |
| Hiseq-MECP2-P7F1 | CAAGCAGAAGACGGCATACGAGATCGTACAGTGACTGGAGTTCAGACGTGTGCTCTTCCGATCTCTGCCCTGTAGAGATAGGAGTTG |
| Hiseq-MECP2-P7F1-a | CAAGCAGAAGACGGCATACGAGATGATCTGGTGACTGGAGTTCAGACGTGTGCTCTTCCGATCTCTGCCCTGTAGAGATAGGAGTTG |
| Hiseq-MECP2-P7R1 | CAAGCAGAAGACGGCATACGAGATGTAGCTGTGACTGGAGTTCAGACGTGTGCTCTTCCGATCTGCAAAGCAGAGACATCAGAAGG |
| Hiseq-MECP2-P7R1-a | CAAGCAGAAGACGGCATACGAGATCTTGTAGTGACTGGAGTTCAGACGTGTGCTCTTCCGATCTGCAAAGCAGAGACATCAGAAGG |
| Hiseq-MECP2-P5F1 | AATGATACGGCGACCACCGAGATCTACACTCTTTCCCTACACGACGCTCTTCCGATCTTCCTCTGAGCGGAAACACTG |
| Hiseq-MECP2-P5F1-a | AATGATACGGCGACCACCGAGATCTACACTCTTTCCCTACACGACGCTCTTCCGATCTAGTTCCTCTGAGCGGAAACACTG |
| Hiseq-MECP2-P5F1-b | AATGATACGGCGACCACCGAGATCTACACTCTTTCCCTACACGACGCTCTTCCGATCTTACGTCCTCTGAGCGGAAACACTG |
| Hiseq-MECP2-P5R1 | AATGATACGGCGACCACCGAGATCTACACTCTTTCCCTACACGACGCTCTTCCGATCTCTCTTGTCAGCCATCGAGCC |
| Hiseq-MECP2-P5R1-a | AATGATACGGCGACCACCGAGATCTACACTCTTTCCCTACACGACGCTCTTCCGATCTACACTCTTGTCAGCCATCGAGCC |
| Hiseq-MECP2-P5R1-b | AATGATACGGCGACCACCGAGATCTACACTCTTTCCCTACACGACGCTCTTCCGATCTTCGACTCTTGTCAGCCATCGAGCC |
| Hiseq-MAZ-P7F1 | CAAGCAGAAGACGGCATACGAGATGTCGAAGTGACTGGAGTTCAGACGTGTGCTCTTCCGATCTACTGTGGCAAGAGCTTCTCC |
| Hiseq-MAZ-P7F1-a | CAAGCAGAAGACGGCATACGAGATAGGAATGTGACTGGAGTTCAGACGTGTGCTCTTCCGATCTACTGTGGCAAGAGCTTCTCC |
| Hiseq-MAZ-P7R1 | CAAGCAGAAGACGGCATACGAGATGCAATCGTGACTGGAGTTCAGACGTGTGCTCTTCCGATCTCTTAAGCAGGAAATCCCTCCCC |
| Hiseq-MAZ-P7R1-a | CAAGCAGAAGACGGCATACGAGATACTTGAGTGACTGGAGTTCAGACGTGTGCTCTTCCGATCTCTTAAGCAGGAAATCCCTCCCC |
| Hiseq-MAZ-P5F1 | AATGATACGGCGACCACCGAGATCTACACTCTTTCCCTACACGACGCTCTTCCGATCTACAAAGGTACATGCCGAGGG |
| Hiseq-MAZ-P5F1-a | AATGATACGGCGACCACCGAGATCTACACTCTTTCCCTACACGACGCTCTTCCGATCTCCGACAAAGGTACATGCCGAGGG |
| Hiseq-MAZ-P5F1-b | AATGATACGGCGACCACCGAGATCTACACTCTTTCCCTACACGACGCTCTTCCGATCTGATCACAAAGGTACATGCCGAGGG |
| Hiseq-MAZ-P5R1 | AATGATACGGCGACCACCGAGATCTACACTCTTTCCCTACACGACGCTCTTCCGATCTTCCCTTGACCTCTTGTAGGAATCT |
| Hiseq-MAZ-P5R1-a | AATGATACGGCGACCACCGAGATCTACACTCTTTCCCTACACGACGCTCTTCCGATCTCACTCCCTTGACCTCTTGTAGGAATCT |
| Hiseq-MAZ-P5R1-b | AATGATACGGCGACCACCGAGATCTACACTCTTTCCCTACACGACG |

CTCTTCCGATCTATGCTCCCTTGACCTCTTGTAGGAATCT

Hiseq-PARP1-P7F1 CAAGCAGAAGACGGCATACGAGATTCAGGTGTGACTGGAGTTCAGA
CGTGTGCTCTTCCGATCTGCTTTATTGAGGCAGCAGTGTTATG

Hiseq-PARP1-P7F1-a CAAGCAGAAGACGGCATACGAGATCACTGTGTGACTGGAGTTCAGA
CGTGTGCTCTTCCGATCTGCTTTATTGAGGCAGCAGTGTTATG

Hiseq-PARP1-P7R1 CAAGCAGAAGACGGCATACGAGATGGAACTGTGACTGGAGTTCAGA
CGTGTGCTCTTCCGATCTACTAACTAAAGCAGGGACAGGG

Hiseq-PARP1-P7R1-a CAAGCAGAAGACGGCATACGAGATATGAGCGTGACTGGAGTTCAGA
CGTGTGCTCTTCCGATCTACTAACTAAAGCAGGGACAGGG

Hiseq-PARP1-P5F1 AATGATACGGCGACCACCGAGATCTACACTCTTTCCCTACACGACG
CTCTTCCGATCTTCTCTTAGGACACCAAACACAGC

Hiseq-PARP1-P5F1-a AATGATACGGCGACCACCGAGATCTACACTCTTTCCCTACACGACG
CTCTTCCGATCTGGCTCTCTTAGGACACCAAACACAGC

Hiseq-PARP1-P5F1-b AATGATACGGCGACCACCGAGATCTACACTCTTTCCCTACACGACG
CTCTTCCGATCTTCTGTCTCTTAGGACACCAAACACAGC

Hiseq-PARP1-P5R1 AATGATACGGCGACCACCGAGATCTACACTCTTTCCCTACACGACG
CTCTTCCGATCTCATGAGGCAGCTCACCACTAA

Hiseq-PARP1-P5R1-a AATGATACGGCGACCACCGAGATCTACACTCTTTCCCTACACGACG
CTCTTCCGATCTAAGCATGAGGCAGCTCACCACTAA

Hiseq-PARP1-P5R1-b AATGATACGGCGACCACCGAGATCTACACTCTTTCCCTACACGACG
CTCTTCCGATCTCTTGCATGAGGCAGCTCACCACTAA

Hiseq-PRDM5-P7F CAAGCAGAAGACGGCATACGAGATTGACCAGTGACTGGAGTTCAGA
CGTGTGCTCTTCCGATCTCCATCACTGGGAAGCACGAA

Hiseq-PRDM5-P7F-a CAAGCAGAAGACGGCATACGAGATACAGTGGTGACTGGAGTTCAGA
CGTGTGCTCTTCCGATCTCCATCACTGGGAAGCACGAA

Hiseq-PRDM5-P7R CAAGCAGAAGACGGCATACGAGATTGCCATGTGACTGGAGTTCAGA
CGTGTGCTCTTCCGATCTTTACCATATCAGTGTTGCTGGACA

Hiseq-PRDM5-P7R-a CAAGCAGAAGACGGCATACGAGATAAGCTAGTGACTGGAGTTCAGA
CGTGTGCTCTTCCGATCTTTACCATATCAGTGTTGCTGGACA

Hiseq-PRDM5-P5F3 AATGATACGGCGACCACCGAGATCTACACTCTTTCCCTACACGACG
CTCTTCCGATCTTTCATGTCTGTATGACTTTGCTGC

Hiseq-PRDM5-P5F3-a AATGATACGGCGACCACCGAGATCTACACTCTTTCCCTACACGACG
CTCTTCCGATCTGCATTCATGTCTGTATGACTTTGCTGC

Hiseq-PRDM5-P5F3-b AATGATACGGCGACCACCGAGATCTACACTCTTTCCCTACACGACG
CTCTTCCGATCTCGGTTTCATGTCTGTATGACTTTGCTGC

Hiseq-PRDM5-P5R3 AATGATACGGCGACCACCGAGATCTACACTCTTTCCCTACACGACG
CTCTTCCGATCTTGACCAGCACATTATTTCTCTCAGA

Hiseq-PRDM5-P5R3-a AATGATACGGCGACCACCGAGATCTACACTCTTTCCCTACACGACG
CTCTTCCGATCTATATGACCAGCACATTATTTCTCTCAGA

Hiseq-PRDM5-P5R3-b AATGATACGGCGACCACCGAGATCTACACTCTTTCCCTACACGACG
CTCTTCCGATCTTGAGTGACCAGCACATTATTTCTCTCAGA

Hiseq-YY1-P5F1 AATGATACGGCGACCACCGAGATCTACACTCTTTCCCTACACGACG
CTCTTCCGATCTAACTCTGGAGCAGAAAGCCTAAT

Hiseq-YY1-P5F1-a AATGATACGGCGACCACCGAGATCTACACTCTTTCCCTACACGACG

| | |
|---|---|
| | CTCTTCCGATCTGCTAACTCTGGAGCAGAAAGCCTAAT |
| Hiseq-YY1-P5F1-b | AATGATACGGCGACCACCGAGATCTACACTCTTTCCCTACACGACG CTCTTCCGATCTAACTAACTCTGGAGCAGAAAGCCTAAT |
| Hiseq-YY1-P5R1 | AATGATACGGCGACCACCGAGATCTACACTCTTTCCCTACACGACG CTCTTCCGATCTGCAACCCACTCTGTTCTTAGAATG |
| Hiseq-YY1-P5R1-a | AATGATACGGCGACCACCGAGATCTACACTCTTTCCCTACACGACG CTCTTCCGATCTCGTGCAACCCACTCTGTTCTTAGAATG |
| Hiseq-YY1-P5R1-b | AATGATACGGCGACCACCGAGATCTACACTCTTTCCCTACACGACG CTCTTCCGATCTTGCAGCAACCCACTCTGTTCTTAGAATG |
| Hiseq-YY1-P7F2 | CAAGCAGAAGACGGCATACGAGATCGATGTGTGACTGGAGTTCAGA CGTGTGCTCTTCCGATCTCTCCTGACCTAAAGTGATCCACC |
| Hiseq-YY1-P7F2-a | CAAGCAGAAGACGGCATACGAGATCGAAACGTGACTGGAGTTCAGA CGTGTGCTCTTCCGATCTCTCCTGACCTAAAGTGATCCACC |
| Hiseq-YY1-P7R2 | CAAGCAGAAGACGGCATACGAGATTTAGGCGTGACTGGAGTTCAGA CGTGTGCTCTTCCGATCTCTCTACTCACACAGCCCCTTC |
| Hiseq-YY1-P7R2-a | CAAGCAGAAGACGGCATACGAGATCAGATCGTGACTGGAGTTCAGA CGTGTGCTCTTCCGATCTCTCTACTCACACAGCCCCTTC |
| Primers for linear amplifications followed by high throughput genome wide translocation sequencing | |
| Nested-HPRT1-E2F-a | AATGATACGGCGACCACCGAGATCTACACTCTTTCCCTACACGACG CTCTTCCGATCTGCTACCATGCTGAGGATTTGGAAAGGG |
| Nested-HPRT1-E2F-b | AATGATACGGCGACCACCGAGATCTACACTCTTTCCCTACACGACG CTCTTCCGATCTGCTGCACTGCTGAGGATTTGGAAAGGGT |
| Nested-HPRT1-E2F-c | AATGATACGGCGACCACCGAGATCTACACTCTTTCCCTACACGACG CTCTTCCGATCTGCTATCGTGCTGAGGATTTGGAAAGGGT |
| Nested-HPRT1-E2F-d | AATGATACGGCGACCACCGAGATCTACACTCTTTCCCTACACGACG CTCTTCCGATCTGATTCATTGCTGAGGATTTGGAAAGGGT |
| P7-A | CAAGCAGAAGACGGCATACGAGATTCGTACGTGACTGGAGTTCAGA CGTGTGCTCTTCCGATCT |
| P7-B | CAAGCAGAAGACGGCATACGAGATTCAGACGTGACTGGAGTTCAGA CGTGTGCTCTTCCGATCT |
| P7-C | CAAGCAGAAGACGGCATACGAGATTCGCATGTGACTGGAGTTCAGA CGTGTGCTCTTCCGATCT |
| P7-D | CAAGCAGAAGACGGCATACGAGATCGAGAGGTGACTGGAGTTCAGA CGTGTGCTCTTCCGATCT |
| Adapter-upper | GACGTGTGCTCTTCCGATCTGNNNNNN |
| Adapter-lower | CAGATCGGAAGAGCACACGTC |
| Bio-HPRT-E2F | TGTTTGTATCCTGTAATGCTCTCA |
| sgRNA sequences for cell growth assay | |
| HPRT1-E2-sgRNA-F | ACCGTTGCTATTTGAACATAAACT |
| HPRT1-E2-sgRNA-R | AAACAGTTTATGTTCAAATAGCAA |
| HPRT1-E3-sgRNA-F | ACCGGTGGAAGTTTAATGACTAAG |
| HPRT1-E3-sgRNA-R | AAACCTTAGTCATTAAACTTCCAC |
| HPRT1-E3-in-F | ACCGGCCCCCCTTGAGCACACAGA |

| | |
|---|---|
| HPRT1-E3-in-R | AAACTCTGTGTGCTCAAGGGGGGC |
| HPRT1-D-E2-F | ACCGAGTATCAGTTGTGGTATAGT |
| HPRT1-D-E2-R | AAACACTATACCACAACTGATACT |
| DCK-E4-D-sgRNA-F | ACCGATATTTAGAACTCTTTTCAG |
| DCK-E4-D-sgRNA-R | AAACCTGAAAAGAGTTCTAAATAT |
| DCK-E5-U-sgRNA-F | ACCGAGAGATGGAAGAAAAAGGCA |
| DCK-E5-U-sgRNA-R | AAACTGCCTTTTTCTTCCATCTCT |
| DCK-E4-in-F | ACCGAGTGGACAATTTATCAAGAC |
| DCK-E4-in-R | AAACGTCTTGATAAATTGTCCACT |
| SgRNA sequences for the induction of chromosomal rearrangements for linear amplifications | |
| HPRT1-sgRNA1-F | ACCGGTAAGTAAGATCTTAAAATG |
| HPRT1-sgRNA1-R | AAACCATTTTAAGATCTTACTTAC |
| HPRT1-sgRNA2-F | ACCGAGTCCTACAGAAATAAAATC |
| HPRT1-sgRNA2-R | AAACGATTTTATTTCTGTAGGACT |

**Supplementary Note S1.** Sequence alignments designed specifically for CRISPR cleavages.

The power of CRISPR in genome editing greatly promotes the development of biology. One of the main tasks of CRISPR analysis is to align reads to the targeting sites and to call for insertions and/or deletions (indels) at the cleavage site. Although several computational tools have been developed to solve them (1-4), the alignment methods of available tools still have at least two unsolved problems. First, the case with deletion followed by insertion (indel) is not considered. Even for the very recent work (1), only simulation data with either deletion or insertion are tested, and emphasis is mainly put on the ability to predict "long" deletion or insertion. The main problem is that traditional global or local aligners used by current tools, say BWA-MEM (5), Novoalign, and the Needleman-Wunsch algorithm (6), often explain indels as mismatches. Second, chromosomal rearrangements by Cas9 programmed with dual sgRNAs have been used to engineer 3D genomes and to investigate structural variations in cancers, but current tools may predict counterintuitive outcomes for CRISPR dual cleavages (7) (8). For example, considering the following alignment given by the traditional method with the expected cleavage site indicated by ":",

TCAGT**CCT**GAC:CT--TTCTAGTGG

TCAGT**CCT**GAC:CTAGTTCTAGTGG

There is a difficulty in explaining the joining of AC and CT. A more realistic alignment is: first delete CT from the reference, and then insert CTAG.

TCAGT**CCT**GAC:----:CTTTCTAGTGG

TCAGT**CCT**GAC:CTAG:--TTCTAGTGG

Note that the second alignment leaves 'CT' unaligned, thereby considered to have a lower score by traditional methods. We propose a new method to solve this kind of problems in Supplementary Note S2.

**Supplementary Note S2.** Dynamic programming of alignment algorithm for CRISPR cleavages.

Let $X$ be the reference sequence and $O$ be the query sequence (e.g., NGS reads). The famous Smith-Waterman local alignment format is

$$E_{s,w} = \begin{cases} -\infty, & w = 0, \\ \max\{E_{s,w-1} + u, G_{s,w-1} + v\}, & w > 0. \end{cases}$$

$$F_{s,w} = \begin{cases} -\infty, & s = 0, \\ \max\{F_{s-1,w} + u, G_{s-1,w} + v\}, & s > 0. \end{cases}$$

$$G_{s,w} = \begin{cases} \max\{E_{s,w}, F_{s,w}, 0\}, & s = 0 \vee w = 0, \\ \max\{G_{s-1,w-1} + \gamma(X_s, O_w), E_{s,w}, F_{s,w}, 0\}, & s > 0 \wedge w > 0. \end{cases}$$

Observe the following iteration:

$$A_w = \max_{0 \le s \le |X|} G_{s,w}.$$

$$C_w = \max\{A_w, C_{w-1}\}.$$

$C_w$ is the score of the optimal local alignment of the first $w$ bases of each NGS read $O$ to the reference sequence $X$. Keep the iterations of $E_{s,w}$ and $F_{s,w}$ and replace the zeros in the iteration of $G_{s,w}$ by $C_w$.

$$G_{s,w} = \begin{cases} \max\{E_{s,w}, F_{s,w}, C_w\}, & s = 0 \vee w = 0, \\ \max\{G_{s-1,w-1} + \gamma(X_s, O_w), E_{s,w}, F_{s,w}, C_w\}, & s > 0 \wedge w > 0. \end{cases}$$

The new $C_w$ now becomes the optimal two-segment local alignment of the first $w$ bases of each NGS read $O$ to the reference sequence $X$. More specifically, two non-overlapping parts of $O$ are aligned to $X$ in order. Such an alignment method is suitable for CRISPR analysis for both single and double cleavages.

**Supplementary Note S3.** Generalization of the alignment algorithm.

Our new alignment method for CRISPR analysis can be generalized in four aspects easily. Firstly, if we iterate $C_w$ for $N$ times, then we will get the optimal $N$-segment local alignment of $O$. Secondly, the reference sequence $X$ can be arbitrarily changed in each iteration, thereby segments of $O$ can be aligned to various reference sequences. Thirdly, one may replace the iteration of $C_w$ by

$$B_w = \begin{cases} -\infty, & w = 0, \\ \max\{A_{w-1} + v, B_{w-1} + u\}, & w > 0. \end{cases}$$

$$C_w = \max\{A_w, B_w\}.$$

This introduces an affine penalty for unaligned parts of $O$ between aligned segments. Finally, one may replace the iterations of $G_{s,w}$ and $A_w$ by

$$G_{s,w} = \begin{cases} \max\{E_{s,w}, F_{s,w}, C_w + g_s\}, & s = 0 \lor w = 0, \\ \max\{G_{s-1,w-1} + \gamma(X_s, O_w), E_{s,w}, F_{s,w}, C_w + g_s\}, & s > 0 \land w > 0. \end{cases}$$

$$A_w = \max_{0 \le s \le |X|} \{G_{s,w} + g_{|X|-s}\}.$$

This introduces a penalty $g_l$ for an unaligned part at the beginning or end of the reference sequence $X$, where $l$ is the length of the unaligned part.

**Supplementary Note S4.** The basic usage of customized computer program and its source code in C++

To use the program, move all files (executable program, input files of NGS reads, reference sequences) into the same folder, change path to that folder, and run the program.
Parameter list
-help: Display help.
-files: Input files of NGS reads separated by comma. If not specified, all files in the current path except those specified by -references will be analyzed in dictionary order. File-wise options are supported (separated by comma). If option is not enough to cover all files of reads, then the last option is used for uncovered files.
-references: Must be specified.
-u: The gap extension score. (default: -2)
-v: The gap opening score. (default: -5)
-s0: The base mismatching score. (default: -3)
-s1: The base matching score. (default: +1)
-alg_types: One of "local_imbed", "imbed", "local", "contain".
-DIVCON: Set divide-and-conquer linear space method to "on" or "off". (default: on)
-SEQ_BATCH: Set the number of reads passed in one time. (default: 300)
-per_thres: Percentage threshold of repeated reads. The same read may repeat many times in the file. Percentage of a read is calculated by (repeated # of the read in the file)/(total # of reads in the file). Reads with percentages less than -per_thres are excluded from analyses.
-ini_alpha: Path to the file used to initialize the distribution of the left ligation ends. If not specified, a uniform distribution over the ligation range will be used. Note that the file must be consistent with the information in reference sequences.
-ini_beta: Similar as -ini_alpha but for the right ligation ends.
-ini_pi: Path to the file used to initialize the distribution of the middle insertions. If not specified, a uniform distribution over all possible middle insertions upper to length specified by -MID_MAX will be used.
-ALIGN_MAX: The maximally reserved number of best alignments for each read (Each read may have several best alignments to the reference). (default: 5)
-MID_MAX: The maximal length of middle insertion. (default: 0)
-thres: The terminal threshold for EM algorithm. (default: 0.000001)
-mode: The method to explain the indel (-mode overlapping or -mode nonoverlapping). (default: nonoverlapping)

-disp_left: Displaying-size left to the rearrangement junctions. (default: 20)
-disp_right: Displaying-size right to the rearrangement junctions. (default: 40)
-disp_num: Number of reads to display. (default: 20)
-THR_MAX: Upper limit of threads used. Only specified once and used for all read files (no file-wise specification). (default: the hardware available threads)

**Computer source code (main.cpp) in C++:**

```
#include <iostream>
#include <fstream>
#include <vector>
#include <cstring>
#include <map>
#include <limits>
#include <tuple>
#include <algorithm>
#include <experimental/filesystem>
#include <queue>
#include <mutex>
#include <future>
#include <condition_variable>
#include <functional>
#include <iomanip>
#include <cmath>
#include <numeric>

struct Command_content
{
	std::vector<std::string> disp_left;
	std::vector<std::string> disp_right;
	std::vector<std::string> disp_num;
	std::vector<std::string> DIVCON;
	std::vector<std::string> SEQ_BATCH;
	std::vector<std::string> ALIGN_MAX;
	std::vector<std::string> S0;
	std::vector<std::string> S1;
	std::vector<std::string> u;
	std::vector<std::string> v;
	std::vector<std::string> references;
	std::vector<std::string> files;
	std::vector<std::string> alg_types;
	std::vector<std::string> per_thres;
	std::vector<std::string> ini_alpha;
	std::vector<std::string> ini_beta;
	std::vector<std::string> ini_pi;
```

```
	std::vector<std::string> MID_MAX;
	std::vector<std::string> thres;
	std::vector<std::string> mode;
	std::vector<std::string> THR_MAX;
};

struct Back
{
	int max_val;
	std::vector<Back*> biters;
};

template<typename T>
class TD_array
{
	std::vector<T> data;
	int row;
	int col;
	int rowcol;
	int slice;
public:
	TD_array(int row_=0, int col_=0, int slice_=0)
	{
		row=row_;
		col=col_;
		rowcol=row*col;
		slice=slice_;
		data.resize(row*col*slice);
	}
	TD_array(int row_, int col_, int slice_, int val_)
	{
		row=row_;
		col=col_;
		rowcol=row*col;
		slice=slice_;
		data.resize(row*col*slice, val_);
	}
	void resize(int row_, int col_, int slice_)
	{
		row=row_;
		col=col_;
		rowcol=row*col;
		slice=slice_;
		data.resize(row*col*slice);
```

```
}
void resize(int row_, int col_, int slice_, int val_)
{
	row=row_;
	col=col_;
	rowcol=row*col;
	slice=slice_;
	data.resize(row*col*slice, val_);
}
T& operator()(int i, int j, int k)
{
	return data[rowcol*k+row*j+i];
}
void fill(T val)
{
	std::fill(data.begin(),data.end(),val);
}
double accumulate()
{
	return accumulate(data.begin(), data.end(), 0.0);
}
int gsli(T* ptr)
{
	return (ptr-data.data())/rowcol;
}
int gcol(T* ptr)
{
	return ((ptr-data.data())%rowcol)/row;
}
int grow(T* ptr)
{
	return (ptr-data.data())%row;
}
std::vector<int>::iterator p2i(int* ptr)
{
	return data.begin()+(ptr-data.data());
}
int size(int dim)
{
	if(dim==0)
		return row;
	else if(dim==1)
		return col;
	else if(dim==2)
```

```
                return slice;
            else
                return -1;
        }
        std::tuple<std::vector<T>,std::vector<T>,std::vector<T>> boundary_sum()
        {
            std::vector<T> alpha(row,0), beta(col,0), pi(slice,0);
            std::vector<T> tmp(col*slice,0);
            int index=0;
            int tind=0;
            for(int s=0;s<slice;++s)
                for(int c=0;c<col;++c)
                {
                    for(int r=0;r<row;++r)
                    {
                        alpha[r]+=data[index];
                        tmp[tind]+=data[index];
                        ++index;
                    }
                    beta[c]+=tmp[tind];
                    pi[s]+=tmp[tind];
                    ++tind;
                }
            return std::make_tuple(alpha, beta, pi);
        }
};

template<typename T>
class thread_safe_queue
{
private:
        mutable std::mutex mut;
        std::queue<T> data_queue;
        std::condition_variable data_cond;
public:
        thread_safe_queue()
        {}
        void push(T new_value)
        {
            std::lock_guard<std::mutex> lk(mut);
            data_queue.push(std::move(new_value));
            data_cond.notify_one();
        }
        bool try_pop(T& value)
```

```
    {
    std::lock_guard<std::mutex> lk(mut);
    if(data_queue.empty())
        return false;
    value=std::move(data_queue.front());
    data_queue.pop();
    return true;
    }
    bool empty() const
    {
        std::lock_guard<std::mutex> lk(mut);
        return data_queue.empty();
    }
};

class join_threads
{
    std::vector<std::thread>& threads;
public:
    explicit join_threads(std::vector<std::thread>& threads_):
        threads(threads_)
    {}
    ~join_threads()
    {
        for(unsigned long i=0;i<threads.size();++i)
            if(threads[i].joinable())
                threads[i].join();
    }
};

class function_wrapper
{
    struct impl_base
    {
        virtual void call()=0;
        virtual ~impl_base() {}
    };
    std::unique_ptr<impl_base> impl;
    template<typename F>
    struct impl_type: impl_base
    {
        F f;
        impl_type(F&& f_): f(std::move(f_)) {}
        void call() { f(); }
```

```
    };
public:
    template<typename F>
    function_wrapper(F&& f):
        impl(new impl_type<F>(std::move(f)))
    {}
    void operator()() { impl->call(); }
    function_wrapper() = default;
    function_wrapper(function_wrapper&& other):
        impl(std::move(other.impl))
    {}
    function_wrapper& operator=(function_wrapper&& other)
    {
        impl=std::move(other.impl);
        return *this;
    }
    function_wrapper(const function_wrapper&)=delete;
    function_wrapper(function_wrapper&)=delete;
    function_wrapper& operator=(const function_wrapper&)=delete;
};

class thread_pool
{
    std::atomic_bool done;
    thread_safe_queue<function_wrapper> work_queue;
    std::vector<std::thread> threads;
    join_threads joiner;
    void worker_thread()
    {
        while(!done)
        {
            function_wrapper task;
            if(work_queue.try_pop(task))
                task();
            else
                std::this_thread::yield();
        }
    }
public:
    thread_pool(int THR_MAX=std::thread::hardware_concurrency()):
        done(false),joiner(threads)
    {
        try
        {
```

```
			for(unsigned i=0;i<THR_MAX;++i)
				threads.push_back(std::thread(&thread_pool::worker_thread,this));
		}
		catch(...)
		{
			done=true;
			throw;
		}
	}
	~thread_pool()
	{
		done=true;
	}
	template<typename FunctionType>
	std::future<typename std::result_of<FunctionType()>::type>
		submit(FunctionType f)
	{
		typedef typename std::result_of<FunctionType()>::type result_type;
		std::packaged_task<result_type()> task(std::move(f));
		std::future<result_type> res(task.get_future());
		work_queue.push(std::move(task));
		return res;
	}
};

struct Align
{
	int index, max_score;
	double num;
	std::vector<int> left, right, c_left, c_right, c_left_in, c_right_in;
	std::vector<std::string> MID, c_MID, c_MID_in;
	std::pair<std::string,std::string> alignment;
};

struct JuncIndel
{
	double num;
	std::vector<std::string> ligation_indel;
	std::vector<std::vector<int>> insert;
};

class MY_LESS
{
	std::string& ref;
```

```
public:
	MY_LESS(std::string& x) : ref(x) {}
	bool operator()(const std::tuple<int,int,std::string>& LRM1, const
std::tuple<int,int,std::string>& LRM2) const
	{
		int i=std::min(std::get<0>(LRM1),std::get<0>(LRM2));
		int up1=ref.size()+std::get<0>(LRM1)+std::get<2>(LRM1).size()-
std::get<1>(LRM1);
		int up2=ref.size()+std::get<0>(LRM2)+std::get<2>(LRM2).size()-
std::get<1>(LRM2);
		while(i<up1 && i<up2)
		{
			char C1, C2;
			if(i<std::get<0>(LRM1))
				C1=ref[i];
			else if(i<std::get<0>(LRM1)+std::get<2>(LRM1).size())
				C1=std::get<2>(LRM1)[i-std::get<0>(LRM1)];
			else
				C1=ref[i-std::get<0>(LRM1)-
std::get<2>(LRM1).size()+std::get<1>(LRM1)];
			if(i<std::get<0>(LRM2))
				C2=ref[i];
			else if(i<std::get<0>(LRM2)+std::get<2>(LRM2).size())
				C2=std::get<2>(LRM2)[i-std::get<0>(LRM2)];
			else
				C2=ref[i-std::get<0>(LRM2)-
std::get<2>(LRM2).size()+std::get<1>(LRM2)];

			if(C1<C2)
				return true;
			else if(C1>C2)
				return false;
			++i;
		}
		if(up1<up2)
			return true;
		else
			return false;
	}
};

std::vector<std::pair<std::vector<int>,std::string>> unify_read(std::string file, int&
total_read);
```

```
std::vector<Align> wapper_column_wise(std::string x, std::vector<int> S, std::string alg_type,
int u, int v, int ALIGN_MAX, std::vector<std::string> os, std::vector<int> index,
std::vector<double> num, int max_len, int S0, int S1, std::string file, std::vector<int> left_exp,
std::vector<int> right_exp, std::string mode);

std::vector<Align> column_wise(std::vector<int>::iterator ve_b, std::vector<int>::iterator
ue_b, int row, std::vector<int>::iterator vf_b, std::vector<int>::iterator uf_b,
std::vector<int>::iterator tvf_b, std::vector<int>::iterator tuf_b, TD_array<int>& gamma,
std::string::iterator x_b, std::string::iterator o_b, std::string::iterator o_e,
std::vector<int>::iterator S_b, std::vector<int>::iterator S_e, bool head, bool tail, bool ceil,
bool floor, int ALIGN_MAX, std::map<char,int>& nt2int, TD_array<Back>& EFG);

void ori_cmp(Back& back, int new_val, Back* address);

std::tuple<std::map<char,int>,TD_array<int>,std::vector<int>,std::vector<int>,std::vector<in
t>,std::vector<int>,TD_array<int>,TD_array<int>> initialization(std::vector<int>& S,
std::string& alg_type, int u, int v, int max_len, int S0, int S1);

std::vector<Align> wapper_divide_and_conquer(std::string x, std::vector<int> S, std::string
alg_type, int u, int v, std::vector<std::string> os, std::vector<int> index, std::vector<double>
num, int max_len, int S0, int S1, std::string file, std::vector<int> left_exp, std::vector<int>
right_exp, std::string mode);

void divide_and_conquer(std::vector<int>::iterator ve_b, std::vector<int>::iterator ue_b, int
row, std::vector<int>::iterator vf_b, std::vector<int>::iterator uf_b, std::vector<int>::iterator
tvf_b, std::vector<int>::iterator tuf_b, TD_array<int>& gamma, std::string::iterator x_b,
std::string::iterator o_b, std::string::iterator o_e, std::vector<int>::iterator S_b,
std::vector<int>::iterator S_e, bool head, bool tail, bool ceil, bool floor, std::map<char,int>&
nt2int, Align& align, std::vector<int>::iterator GU_b, std::vector<int>::reverse_iterator GD_b,
std::vector<int>::iterator GL_b, std::vector<int>::reverse_iterator GR_b,
std::vector<int>::iterator EL_b, std::vector<int>::reverse_iterator ER_b);

template<typename vi, typename si>
void row_wise_linear(vi ve_b, vi ue_b, int row, vi vf_b, vi uf_b, vi tvf_b, vi tuf_b,
TD_array<int>& gamma, si x_b, si o_b, si o_e, vi S_b, vi S_e, std::map<char,int>& nt2int, vi
G_b);

template<typename vi, typename si>
void column_wise_linear(vi ve_b, vi ue_b, int row, vi vf_b, vi uf_b, vi tvf_b, vi tuf_b,
TD_array<int>& gamma, si x_b, si o_b, si o_e, vi S_b, vi S_e, bool head, bool ceil, bool floor,
std::map<char,int>& nt2int, vi E_b, vi G_b);
```

```
void load_ref(std::string& ref_file, std::string& x, std::vector<int>& S, std::vector<int>&
left_exp, std::vector<int>& left_down, std::vector<int>& left_up, std::vector<int>& right_exp,
std::vector<int>& right_down, std::vector<int>& right_up);
std::tuple<std::vector<std::string>,std::vector<int>,std::vector<double>>
load_read(std::vector<std::pair<std::vector<int>,std::string>>& sus_ord, int& so, int&
hasgot, int& max_len, int SEQ_BATCH, double per_thres, int total_read, std::ofstream& fout);
std::vector<std::string> my_split(std::string line, std::string sc);
Command_content command(int argc, char **argv);
void getFiles(std::vector<std::string> & files, std::vector<std::string> & references);

std::tuple<std::vector<int>,std::vector<int>>
extract_to_pair(std::pair<std::string,std::string>& alignment, std::vector<int>& left,
std::vector<int>& right);
std::tuple<std::vector<int>,std::vector<int>> normal_2_c(std::vector<int>& left,
std::vector<int>& right, std::vector<int>& left_exp, std::vector<int>& right_exp, std::string&
mode);
std::vector<std::string> get_mid(std::pair<std::string,std::string>& alignment,
std::vector<int>& left, std::vector<int>& right);
std::tuple<std::vector<int>,std::vector<int>,std::vector<std::string>,std::vector<int>,std::vect
or<int>,std::vector<std::string>,std::vector<int>,std::vector<int>,std::vector<std::string>>
search_left_right(std::pair<std::string,std::string>& alignment, std::vector<int>& S,
std::vector<int>& left_exp, std::vector<int>& right_exp, std::string& mode, std::string& x);

void disp_sample(std::vector<Align>& aligns, std::vector<int>& left_exp, std::vector<int>&
right_exp, int disp_left, int disp_right, int disp_num, std::string& x, std::string& mode,
std::string& file, int total_read);

void indel_label_fun(std::vector<Align>& aligns,
std::vector<std::pair<std::vector<int>,std::string>>& sus_ord, int total_read,
std::vector<int>& left_exp, std::vector<int>& right_exp, std::string& file);

void div_num_fun(std::vector<Align>::iterator first, std::vector<Align>::iterator last);

void EM_predict(std::vector<Align>& aligns, std::string& x, std::vector<int>& left_exp,
std::vector<int>& left_down, std::vector<int>& left_up, std::vector<int>& right_exp,
std::vector<int>& right_down, std::vector<int>& right_up, int MID_MAX, std::string&
ini_alpha, std::string& ini_beta, std::string& ini_pi, double thres, std::string& file);

std::vector<std::vector<double>> initial_EM(std::string& ini, int dim,
std::vector<TD_array<double>>& N_mats);

int main(int argc, char **argv)
{
	for(int i=1; i<argc; ++i)
```

```
if(!strcmp(argv[i],"-help"))
{
	std::cout << "Basic usage\n"
		<< "To use the program, move all files (executable program, input files of NGS reads, reference sequences) into the same folder, change path to that folder, and run the program\n"
		<< "Parameter list\n"
		<< "-help: Display help.\n"
		<< "-files: Input files of NGS reads separated by comma. If not specified, all files in the current path except those specified by -references will be analyzed in dictionary order. File-wise options are supported (separated by comma). If option is not enough to cover all files of reads, then the last option is used for uncovered files.\n"
		<< "-references: Must be specified.\n"
		<< "-u: The gap extension score. (default: -2)\n"
		<< "-v: The gap opening score. (default: -5)\n"
		<< "-s0: The base mismatching score. (default: -3)\n"
		<< "-s1: The base matching score. (default: +1)\n"
		<< "-alg_types: One of \"local_imbed\", \"imbed\", \"local\", \"contain\".\n"
		<< "-DIVCON: Set divide-and-conquer linear space method to \"on\" or \"off\". (default: on)\n"
		<< "-SEQ_BATCH: Set the number of reads passed in one time. (default: 300)\n"
		<< "-per_thres: Percentage threshold of repeated reads. The same read may repeat many times in the file. Percentage of a read is calculated by (repeated # of the read in the file)/(total # of reads in the file). Reads with percentages less than -per_thres are excluded from analyses.\n"
		<< "-ini_alpha: Path to the file used to initialize the distribution of the left ligation ends. If not specified, a uniform distribution over the ligation range will be used. Note that the file must be consistent with the information in reference sequences.\n"
		<< "-ini_beta: Similar as -ini_alpha but for the right ligation ends.\n"
		<< "-ini_pi: Path to the file used to initialize the distribution of the middle insertions. If not specified, a uniform distribution over all possible middle insertions upper to length specified by -MID_MAX will be used.\n"
		<< "-ALIGN_MAX: The maximally reserved number of best alignments for each read (Each read may have several best alignments to the reference). (default: 5)\n"
		<< "-MID_MAX: The maximal length of middle insertion. (default: 0)\n"
		<< "-thres: The terminal threshold for EM algorithm. (default: 0.000001)\n"
		<< "-mode: The method to explain the indel (-mode overlapping or -mode nonoverlapping). (default: nonoverlapping)\n"
		<< "-disp_left: Displaying-size left to the rearrangement junctions. (default: 20)\n"
		<< "-disp_right: Displaying-size right to the rearrangement junctions. (default: 40)\n"
		<< "-disp_num: Number of reads to display. (default: 20)\n"
```

```
			<< "-THR_MAX: Upper limit of threads used. Only specified once and used
for all read files (no file-wise specification). (default: the hardware available threads)\n";
			return 0;
		}

	Command_content cc=command(argc, argv);
	if (!cc.files.empty())
	{
		std::ofstream fout("draw.sh");
		fout << "for file in *.gp; do\n\tgnuplot $file;\ndone\n";
		fout.close();
		fout.open("draw.bat");
		fout << "for %%i in (*.gp) do %%i";
		fout.close();
	}
	thread_pool threads(stoi(cc.THR_MAX[0]));

	for(int fi=0; fi<cc.files.size(); ++fi)
	{
		std::string x;
		std::vector<int> S, left_exp, left_down, left_up, right_exp, right_down, right_up;
		load_ref(cc.references[fi], x, S, left_exp, left_down, left_up, right_exp, right_down,
right_up);
		if(x.empty()) return -1;

		int total_read;
		std::vector<std::pair<std::vector<int>,std::string>> sus_ord=unify_read(cc.files[fi],
total_read);
		std::ofstream fout(cc.files[fi]+".sus");
		int so=0, hasgot, max_len;
		std::vector<std::future<std::vector<Align>>> futures;
		while(so<sus_ord.size() &&
double(sus_ord[so].first.size())/double(total_read)>=stod(cc.per_thres[fi]))
		{
			std::vector<std::string> os;
			std::vector<int> index;
			std::vector<double> num;
			std::tie(os,index,num)=load_read(sus_ord, so, hasgot, max_len,
stoi(cc.SEQ_BATCH[fi]), stod(cc.per_thres[fi]), total_read, fout);
			if(stoi(cc.ALIGN_MAX[fi])>1 || cc.DIVCON[fi]!="on")
				futures.push_back(threads.submit(std::bind(wapper_column_wise, x, S,
cc.alg_types[fi], stoi(cc.u[fi]), stoi(cc.v[fi]), stoi(cc.ALIGN_MAX[fi]), std::move(os),
std::move(index), std::move(num), max_len, stoi(cc.S0[fi]), stoi(cc.S1[fi]), cc.files[fi], left_exp,
right_exp, cc.mode[fi])));
```

```
			else
				futures.push_back(threads.submit(std::bind(wapper_divide_and_conquer,
x, S, cc.alg_types[fi], stoi(cc.u[fi]), stoi(cc.v[fi]), std::move(os), std::move(index),
std::move(num), max_len, stoi(cc.S0[fi]), stoi(cc.S1[fi]), cc.files[fi], left_exp, right_exp,
cc.mode[fi])));
		}
		fout.close();
		std::vector<Align> aligns;
		for(unsigned i=0; i<futures.size(); i++)
		{
			std::vector<Align> tmp=futures[i].get();
			std::move(tmp.begin(), tmp.end(), std::back_inserter(aligns));
		}
		fout.open(cc.files[fi]+".alg");
		for(int i=0; i<aligns.size(); ++i)
		{
			fout << aligns[i].index << '\t' << aligns[i].num << '\t' << aligns[i].max_score
<< '\t';
			for(int j=0; j<aligns[i].left.size(); ++j)
				fout << aligns[i].MID[j] << '\t' << aligns[i].right[j] << '\t' << aligns[i].left[j]
<< '\t';
			fout << aligns[i].MID.back() << '\t';
			for(int j=0; j<aligns[i].c_left.size(); ++j)
				fout << aligns[i].c_MID[j] << '\t' << aligns[i].c_right[j] << '\t' <<
aligns[i].c_left[j] << '\t';
			fout << aligns[i].c_MID.back() << '\t';
			for(int j=0; j<aligns[i].c_left_in.size(); ++j)
				fout << aligns[i].c_MID_in[j] << '\t' << aligns[i].c_right_in[j] << '\t' <<
aligns[i].c_left_in[j] << '\t';
			fout << aligns[i].c_MID_in.back() << '\n' << aligns[i].alignment.first << '\n' <<
aligns[i].alignment.second << '\n';
		}
		fout.close();
		disp_sample(aligns, left_exp, right_exp, stoi(cc.disp_left[fi]), stoi(cc.disp_right[fi]),
stoi(cc.disp_num[fi]), x, cc.mode[fi], cc.files[fi], total_read);
		indel_label_fun(aligns, sus_ord, total_read, left_exp, right_exp, cc.files[fi]);
		EM_predict(aligns, x, left_exp, left_down, left_up, right_exp, right_down, right_up,
stoi(cc.MID_MAX[fi]), cc.ini_alpha[fi], cc.ini_beta[fi], cc.ini_pi[fi], stod(cc.thres[fi]), cc.files[fi]);
	}
	return 0;
}

std::vector<std::pair<std::vector<int>,std::string>> unify_read(std::string file, int& total_read)
{
```

```
	std::ifstream fin(file);
	std::map<std::string,std::vector<int>> sus;
	total_read=0;
	std::string str_tmp;
	fin >> str_tmp;
	while(fin.good())
	{
		bool DNA=true;
		for(std::string::iterator iter=str_tmp.begin(); iter!=str_tmp.end(); ++iter)
		{
			*iter=toupper(*iter);
			if(*iter!='N' && *iter!='A' && *iter!='C' && *iter!='G' && *iter!='T')
			{
				DNA=false;
				break;
			}
		}
		if(DNA)
		{
			sus[str_tmp].push_back(total_read);
			total_read++;
		}
		fin >> str_tmp;
	}
	fin.close();
	auto vec_len_more=[](std::pair<std::vector<int>,std::string> & pa1,
std::pair<std::vector<int>,std::string> & pa2){return pa1.first.size()>pa2.first.size();};
	std::vector<std::pair<std::vector<int>,std::string>> sus_ord;
	for(std::map<std::string,std::vector<int>>::iterator iter=sus.begin(); iter!=sus.end();
++iter)
		sus_ord.push_back(std::make_pair(iter->second,iter->first));
	std::sort(sus_ord.begin(), sus_ord.end(), vec_len_more);
	return sus_ord;
}

std::vector<Align> wapper_column_wise(std::string x, std::vector<int> S, std::string alg_type,
int u, int v, int ALIGN_MAX, std::vector<std::string> os, std::vector<int> index,
std::vector<double> num, int max_len, int S0, int S1, std::string file, std::vector<int> left_exp,
std::vector<int> right_exp, std::string mode)
{
	std::map<char,int> nt2int;
	TD_array<int> gamma, vf, uf;
	std::vector<int> ve, ue, tvf, tuf;
```

```
    std::tie(nt2int,gamma,ve, ue, tvf, tuf, vf, uf)=initialization(S, alg_type, u, v, max_len, S0,
S1);
    TD_array<Back> EFG(S.back()+1,max_len+1,3);
    std::vector<Align> aligns;
    for(int i=0; i<os.size(); ++i)
    {
        std::vector<Align> tmp=column_wise(ve.begin(), ue.begin(), S.size()-1,
vf.p2i(&vf(0,0,0)), uf.p2i(&uf(0,0,0)), tvf.begin(), tuf.begin(), gamma, x.begin(), os[i].begin(),
os[i].end(), S.begin(), S.end()-1, true, true, true, true, ALIGN_MAX, nt2int, EFG);
        for(int j=0; j<tmp.size(); ++j)
        {
            tmp[j].index=index[i];
            tmp[j].num=num[i];
            std::tie(tmp[j].left, tmp[j].right, tmp[j].MID, tmp[j].c_left, tmp[j].c_right,
tmp[j].c_MID, tmp[j].c_left_in, tmp[j].c_right_in,
tmp[j].c_MID_in)=search_left_right(tmp[j].alignment, S, left_exp, right_exp, mode, x);
        }
        std::move(tmp.begin(), tmp.end(), std::back_inserter(aligns));
    }
    return aligns;
}

std::tuple<std::map<char,int>,TD_array<int>,std::vector<int>,std::vector<int>,std::vector<in
t>,std::vector<int>,TD_array<int>,TD_array<int>> initialization(std::vector<int>& S,
std::string& alg_type, int u, int v, int max_len, int S0, int S1)
{
    TD_array<int> gamma(5,5,1,S0);
    gamma(1,1,0)=gamma(2,2,0)=gamma(3,3,0)=gamma(4,4,0)=S1;
    std::vector<int> ve(S.back()+1,v), ue(S.back()+1,u), tvf(S.size()-1,v), tuf(S.size()-1,u);
    for(int j=0; j<S.size(); ++j)
    {
        if(alg_type=="local" || alg_type=="contain")
        {
            ve[S[j]]=0;
            ue[S[j]]=0;
        }
        if(alg_type=="local_imbed")
        {
            if(j!=0 && j!=S.size()-1)
            {
                ve[S[j]]=0;
                ue[S[j]]=0;
            }
        }
```

```
			if(alg_type=="local" || alg_type=="local_imbed" || alg_type=="imbed")
			{
				if(j!=S.size()-1)
				{
					tvf[j]=0;
					tuf[j]=0;
				}
			}
		}
		return
std::make_tuple(std::map<char,int>{{'N',0},{'A',1},{'C',2},{'G',3},{'T',4},{'n',0},{'a',1},{'c',2},{'g',3},{'t
',4}},gamma,ve,ue,tvf,tuf,TD_array<int>(S.size()-1,max_len+1,1,v),TD_array<int>(S.size()-
1,max_len+1,1,u));
}

std::vector<Align> column_wise(std::vector<int>::iterator ve_b, std::vector<int>::iterator
ue_b, int row, std::vector<int>::iterator vf_b, std::vector<int>::iterator uf_b,
std::vector<int>::iterator tvf_b, std::vector<int>::iterator tuf_b, TD_array<int>& gamma,
std::string::iterator x_b, std::string::iterator o_b, std::string::iterator o_e,
std::vector<int>::iterator S_b, std::vector<int>::iterator S_e, bool head, bool tail, bool ceil,
bool floor, int ALIGN_MAX, std::map<char,int>& nt2int, TD_array<Back>& EFG)
{
	EFG(0,0,2).max_val=0;
	for(int j=0; j<S_e-S_b; ++j)
		for(int s=*(S_b+j)+1; s<=*(S_b+j+1); ++s)
		{
			if(ceil)
				EFG(s,0,2).max_val=EFG(s-
1,0,2).max_val+((s==*(S_b+j)+1)?(*(tvf_b+j)):(*(tuf_b+j)));
			else
				EFG(s,0,2).max_val=EFG(s-
1,0,2).max_val+((s==*(S_b+j)+1)?(*(vf_b+j)):(*(uf_b+j)));
			EFG(s,0,0).max_val=std::numeric_limits<int>::min()/2;
		}
	for(int w=1; w<=o_e-o_b; ++w)
	{
		EFG(0,w,0).max_val=EFG(0,w,2).max_val=EFG(0,w-1,2).max_val+((w==1 &&
head)?(*ve_b):(*ue_b));
		for(int j=0; j<S_e-S_b; ++j)
		{
			if(floor)
				EFG(*(S_b+j+1),w,1).max_val=std::numeric_limits<int>::min()/2;
			for(int s=*(S_b+j)+1; s<=*(S_b+j+1); ++s)
			{
```

```
                if(floor)
                    ori_cmp(EFG(*(S_b+j+1),w,1), EFG(s-
1,w,2).max_val+*(tvf_b+j)+(*(S_b+j+1)-s)*(*(tuf_b+j)), &EFG(s-1,w,2));
                if(s!=*(S_b+j+1) || !floor)
                {
                    EFG(s,w,1).max_val=std::numeric_limits<int>::min()/2;
                    ori_cmp(EFG(s,w,1), EFG(s-1,w,2).max_val+*(vf_b+j+w*row), &EFG(s-
1,w,2));
                    if(s!=*(S_b+j)+1)
                        ori_cmp(EFG(s,w,1), EFG(s-1,w,1).max_val+*(uf_b+j+w*row),
&EFG(s-1,w,1));
                    if(ceil)
                        ori_cmp(EFG(s,w,1), EFG(*(S_b+j),w,2).max_val+*(tvf_b+j)+(s-
*(S_b+j)-1)*(*(tuf_b+j)), &EFG(*(S_b+j),w,2));
                }
                EFG(s,w,0).max_val=std::numeric_limits<int>::min()/2;
                if(!tail && s==*S_e)
                    ori_cmp(EFG(s,w,0), EFG(s,w-1,2).max_val+*(ue_b+s), &EFG(s,w-1,2));
                else
                {
                    ori_cmp(EFG(s,w,0), EFG(s,w-1,2).max_val+*(ve_b+s), &EFG(s,w-1,2));
                    if(EFG(s,w-1,2).max_val!=EFG(s,w-1,0).max_val ||
*(ve_b+s)!=*(ue_b+s))
                        ori_cmp(EFG(s,w,0), EFG(s,w-1,0).max_val+*(ue_b+s), &EFG(s,w-
1,0));
                }
                EFG(s,w,2).max_val=std::numeric_limits<int>::min()/2;
                ori_cmp(EFG(s,w,2), EFG(s-1,w-1,2).max_val+gamma(nt2int[*(x_b+s-
1)],nt2int[*(o_b+w-1)],0), &EFG(s-1,w-1,2));
                ori_cmp(EFG(s,w,2), EFG(s,w,1).max_val, &EFG(s,w,1));
                ori_cmp(EFG(s,w,2), EFG(s,w,0).max_val, &EFG(s,w,0));
            }
        }
    }
    std::vector<std::vector<std::pair<int,int>>> rivets;
    std::vector<Back*> biters;
    rivets.push_back(std::vector<std::pair<int,int>>());
    biters.push_back(&EFG(*S_e,o_e-o_b,2));
    int i=0;
    while(i<biters.size())
    {
        rivets[i].push_back(std::make_pair(EFG.grow(biters[i]),EFG.gcol(biters[i])));
        if(rivets[i].back().first==0 || rivets[i].back().second==0)
        {
```

```
				rivets[i].push_back(std::make_pair(0,0));
				++i;
			}
			else
			{
				std::vector<Back*>::iterator iter=biters[i]->biters.begin();
				std::vector<Back*>::iterator end_iter=biters[i]->biters.end();
				biters[i]=*iter;
				++iter;
				while(biters.size()<ALIGN_MAX && iter!=end_iter)
				{
					rivets.push_back(rivets[i]);
					biters.push_back(*iter);
					++iter;
				}
			}
		}
		std::vector<Align> aligns;
		for(int i=0; i<rivets.size(); ++i)
		{
			aligns.push_back(Align());
			aligns.back().max_score=EFG(*S_e,o_e-o_b,2).max_val;
			for(std::vector<std::pair<int,int>>::reverse_iterator iter=rivets[i].rbegin();
iter!=rivets[i].rend()-1; ++iter)
			{
				if(iter->first==(iter+1)->first)
				{
					aligns.back().alignment.first.append(std::string((iter+1)->second-
iter->second,'-'));

aligns.back().alignment.second.append(o_b+iter->second,o_b+(iter+1)->second);
				}
				else if(iter->second==(iter+1)->second)
				{
					aligns.back().alignment.first.append(x_b+iter->first,x_b+(iter+1)->first);
					aligns.back().alignment.second.append(std::string((iter+1)->first-
iter->first,'-'));
				}
				else
				{
					aligns.back().alignment.first.append(x_b+iter->first,x_b+(iter+1)->first);

aligns.back().alignment.second.append(o_b+iter->second,o_b+(iter+1)->second);
				}
```

```
		}
	}
	return aligns;
}

void ori_cmp(Back& back, int new_val, Back* address)
{
	if(back.max_val<=new_val)
	{
		if(back.max_val<new_val)
		{
			back.max_val=new_val;
			back.biters.clear();
		}
		back.biters.push_back(address);
	}
}

std::vector<Align> wapper_divide_and_conquer(std::string x, std::vector<int> S, std::string alg_type, int u, int v, std::vector<std::string> os, std::vector<int> index, std::vector<double> num, int max_len, int S0, int S1, std::string file, std::vector<int> left_exp, std::vector<int> right_exp, std::string mode)
{
	std::map<char,int> nt2int;
	TD_array<int> gamma, vf, uf;
	std::vector<int> ve, ue, tvf, tuf;
	std::tie(nt2int,gamma,ve, ue, tvf, tuf, vf, uf)=initialization(S, alg_type, u, v, max_len, S0, S1);
	std::vector<int> GU(max_len+1), GD(max_len+1), GL(S.back()+1), GR(S.back()+1), EL(S.back()+1), ER(S.back()+1);
	std::vector<Align> aligns;
	for(int i=0; i<os.size(); ++i)
	{
		aligns.push_back(Align());
		divide_and_conquer(ve.begin(), ue.begin(), S.size()-1, vf.p2i(&vf(0,0,0)), uf.p2i(&uf(0,0,0)), tvf.begin(), tuf.begin(), gamma, x.begin(), os[i].begin(), os[i].end(), S.begin(), S.end()-1, true, true, true, true, nt2int, aligns.back(), GU.begin(), GD.rbegin(), GL.begin(), GR.rbegin(), EL.begin(), ER.rbegin());
		aligns.back().index=index[i];
		aligns.back().num=num[i];
		std::tie(aligns.back().left, aligns.back().right, aligns.back().MID, aligns.back().c_left, aligns.back().c_right, aligns.back().c_MID, aligns.back().c_left_in, aligns.back().c_right_in, aligns.back().c_MID_in)=search_left_right(aligns.back().alignment, S, left_exp, right_exp, mode, x);
```

```
	}
	return aligns;
}

void divide_and_conquer(std::vector<int>::iterator ve_b, std::vector<int>::iterator ue_b, int
row, std::vector<int>::iterator vf_b, std::vector<int>::iterator uf_b, std::vector<int>::iterator
tvf_b, std::vector<int>::iterator tuf_b, TD_array<int>& gamma, std::string::iterator x_b,
std::string::iterator o_b, std::string::iterator o_e, std::vector<int>::iterator S_b,
std::vector<int>::iterator S_e, bool head, bool tail, bool ceil, bool floor, std::map<char,int>&
nt2int, Align& align, std::vector<int>::iterator GU_b, std::vector<int>::reverse_iterator GD_b,
std::vector<int>::iterator GL_b, std::vector<int>::reverse_iterator GR_b,
std::vector<int>::iterator EL_b, std::vector<int>::reverse_iterator ER_b)
{
	if(o_b==o_e)
	{
		align.alignment.first.append(x_b,x_b+*S_e);
		align.alignment.second.append(std::string(*S_e,'-'));
		for(int i=0; i<S_e-S_b; ++i)
		{
			if(ceil || floor)
				align.max_score+=(*(S_b+i+1)-1)*(*(tuf_b+i))+*(tvf_b+i);
			else
				align.max_score+=(*(S_b+i+1)-1)*(*(uf_b+i))+*(vf_b+i);
		}
		return;
	}
	if(*S_e==0)
	{
		align.alignment.first.append(std::string(o_e-o_b,'-'));
		align.alignment.second.append(o_b,o_e);
		align.max_score+=(o_e-o_b)*(*ue_b);
		if(head || tail)
			align.max_score+=*ve_b-*ue_b;
		return;
	}
	if(o_e-o_b==1)
	{
		TD_array<Back> EFG(*S_e+1,o_e-o_b+1,3);
		std::vector<Align> aligns=column_wise(ve_b, ue_b, row, vf_b, uf_b, tvf_b, tuf_b,
gamma, x_b, o_b, o_e, S_b, S_e, head, tail, ceil, floor, 1, nt2int, EFG);
		align.alignment.first.append(aligns.front().alignment.first);
		align.alignment.second.append(aligns.front().alignment.second);
		align.max_score+=aligns.front().max_score;
		return;
```

```
    }
    if(S_e-S_b>1)
    {
        int js=(S_e-S_b)/2;
        row_wise_linear(ve_b, ue_b, row, vf_b, uf_b, tvf_b, tuf_b, gamma, x_b, o_b, o_e, S_b,
S_b+js, nt2int, GU_b);
        std::vector<int> S_tmp(S_b+js,S_e+1);
        for(std::vector<int>::iterator iter=S_tmp.begin(); iter!=S_tmp.end(); ++iter)
            *iter=*S_e-*iter;
        row_wise_linear(std::vector<int>::reverse_iterator(ve_b+*S_e+1),
std::vector<int>::reverse_iterator(ue_b+*S_e+1), row,
std::vector<int>::reverse_iterator(vf_b+(S_e-S_b)+(o_e-o_b)*row),
std::vector<int>::reverse_iterator(uf_b+(S_e-S_b)+(o_e-o_b)*row),
std::vector<int>::reverse_iterator(tvf_b+(S_e-S_b)),
std::vector<int>::reverse_iterator(tuf_b+(S_e-S_b)), gamma,
std::string::reverse_iterator(x_b+*S_e), std::string::reverse_iterator(o_e),
std::string::reverse_iterator(o_b), S_tmp.rbegin(), S_tmp.rend()-1, nt2int, GD_b);
        int max_tmp=std::numeric_limits<int>::min();
        int ws;
        for(int w=0; w<=o_e-o_b; ++w)
        {
            int tmp=*(GU_b+w)+*(GD_b+(o_e-o_b)-w);
            if(tmp>max_tmp)
            {
                max_tmp=tmp;
                ws=w;
            }
        }
        divide_and_conquer(ve_b, ue_b, row, vf_b, uf_b, tvf_b, tuf_b, gamma, x_b, o_b,
o_b+ws, S_b, S_b+js, true, true, true, true, nt2int, align, GU_b, GD_b, GL_b, GR_b, EL_b,
ER_b);
        for(int i=0; i<S_tmp.size(); ++i)
            S_tmp[i]=*(S_b+js+i)-*(S_b+js);
        divide_and_conquer(ve_b+*(S_b+js), ue_b+*(S_b+js), row, vf_b+js+ws*row,
uf_b+js+ws*row, tvf_b+js, tuf_b+js, gamma, x_b+*(S_b+js), o_b+ws, o_e, S_tmp.begin(),
S_tmp.end()-1, true, true, true, true, nt2int, align, GU_b, GD_b, GL_b, GR_b, EL_b, ER_b);
    }
    else
    {
        int ws=(o_e-o_b)/2;
        column_wise_linear(ve_b, ue_b, row, vf_b, uf_b, tvf_b, tuf_b, gamma, x_b, o_b,
o_b+ws, S_b, S_e, head, ceil, floor, nt2int, EL_b, GL_b);
        std::vector<int> S_tmp{*S_e,*S_b};
```

```
        column_wise_linear(std::vector<int>::reverse_iterator(ve_b+*S_e+1),
std::vector<int>::reverse_iterator(ue_b+*S_e+1), row,
std::vector<int>::reverse_iterator(vf_b+(S_e-S_b)+(o_e-o_b)*row),
std::vector<int>::reverse_iterator(uf_b+(S_e-S_b)+(o_e-o_b)*row),
std::vector<int>::reverse_iterator(tvf_b+(S_e-S_b)),
std::vector<int>::reverse_iterator(tuf_b+(S_e-S_b)), gamma,
std::string::reverse_iterator(x_b+*S_e), std::string::reverse_iterator(o_e),
std::string::reverse_iterator(o_b+ws), S_tmp.rbegin(), S_tmp.rend()-1, tail, floor, ceil, nt2int,
ER_b, GR_b);
        int max_tmp_e=std::numeric_limits<int>::min();
        int max_tmp_g=std::numeric_limits<int>::min();
        int se, sg;
        for(int s=0; s<=*S_e; ++s)
        {
            int tmp_e=*(EL_b+s)+*(ER_b+*S_e-s)-*(ve_b+s)+*(ue_b+s);
            int tmp_g=*(GL_b+s)+*(GR_b+*S_e-s);
            if(tmp_e>max_tmp_e)
            {
                max_tmp_e=tmp_e;
                se=s;
            }
            if(tmp_g>max_tmp_g)
            {
                max_tmp_g=tmp_g;
                sg=s;
            }
        }
        int ss, deltaws;
        bool bool_tmp;
        if(max_tmp_e>max_tmp_g)
        {
            ss=se;
            deltaws=1;
            bool_tmp=false;
        }
        else
        {
            ss=sg;
            deltaws=0;
            bool_tmp=true;
        }
        bool bool2_tmp;
        if(ss==0)
            bool2_tmp=ceil;
```

```
            else if(ss==*S_e)
                  bool2_tmp=floor;
            else
                  bool2_tmp=false;
            S_tmp[0]=0;
            S_tmp[1]=ss;
            divide_and_conquer(ve_b, ue_b, row, vf_b, uf_b, tvf_b, tuf_b, gamma, x_b, o_b,
o_b+ws-deltaws, S_tmp.begin(), S_tmp.end()-1, head, bool_tmp, ceil, bool2_tmp, nt2int,
align, GU_b, GD_b, GL_b, GR_b, EL_b, ER_b);
            align.alignment.first.append(std::string(2*deltaws,'-'));
            align.alignment.second.append(o_b+ws-deltaws,o_b+ws+deltaws);
            if(deltaws>0)
                  align.max_score+=(2*deltaws-1)*(*(ue_b+ss))+*(ve_b+ss);
            S_tmp[0]=0;
            S_tmp[1]=*S_e-ss;
            divide_and_conquer(ve_b+ss, ue_b+ss, row, vf_b+ws+deltaws, uf_b+ws+deltaws,
tvf_b, tuf_b, gamma, x_b+ss, o_b+ws+deltaws, o_e, S_tmp.begin(), S_tmp.end()-1, bool_tmp,
tail, bool2_tmp, floor, nt2int, align, GU_b, GD_b, GL_b, GR_b, EL_b, ER_b);
      }
}

template<typename vi, typename si>
void row_wise_linear(vi ve_b, vi ue_b, int row, vi vf_b, vi uf_b, vi tvf_b, vi tuf_b,
TD_array<int>& gamma, si x_b, si o_b, si o_e, vi S_b, vi S_e, std::map<char,int>& nt2int, vi
G_b)
{
      std::vector<int> tildeG(o_e-o_b+1);
      *G_b=0;
      for(int w=1; w<=o_e-o_b; ++w)
            tildeG[w]=*(G_b+w)=*(G_b+w-1)+((w==1)?(*ve_b):(*ue_b));
      for(int j=0; j<S_e-S_b; ++j)
      {
            std::vector<int> F(o_e-o_b+1);
            std::vector<int> tildeF(o_e-o_b+1,std::numeric_limits<int>::min()/2);
            for(int s=*(S_b+j)+1; s<=*(S_b+j+1); ++s)
            {
                  int hatG=*G_b;
                  *G_b=*G_b+((s==*(S_b+j)+1)?(*(tvf_b+j)):(*(tuf_b+j)));
                  int E=std::numeric_limits<int>::min()/2;
                  for(int w=1; w<=o_e-o_b; ++w)
                  {
                        tildeF[w]=std::max(tildeF[w],*(G_b+w)+*(tvf_b+j)+(*(S_b+j+1)-
s)*(*(tuf_b+j)));
                        E=std::max(*(G_b+w-1)+*(ve_b+s),E+*(ue_b+s));
```

```
                int Gp=*(G_b+w);
                *(G_b+w)=hatG+gamma(nt2int[*(x_b+s-1)],nt2int[*(o_b+w-1)],0);
                *(G_b+w)=std::max(*(G_b+w),E);
                if(s!=*(S_b+j+1))
                {
                    if(s!=*(S_b+j)+1)
                        F[w]=std::max(F[w]+*(uf_b+j+w*row),Gp+*(vf_b+j+w*row));
                    else
                        F[w]=Gp+*(vf_b+j+w*row);
                    F[w]=std::max(F[w],tildeG[w]+*(tvf_b+j)+(s-*(S_b+j)-1)*(*(tuf_b+j)));
                    *(G_b+w)=std::max(*(G_b+w),F[w]);
                }
                else
                {
                    *(G_b+w)=std::max(*(G_b+w),tildeF[w]);
                    tildeG[w]=*(G_b+w);
                }
                hatG=Gp;
            }
        }
    }
}

template<typename vi, typename si>
void column_wise_linear(vi ve_b, vi ue_b, int row, vi vf_b, vi uf_b, vi tvf_b, vi tuf_b,
TD_array<int>& gamma, si x_b, si o_b, si o_e, vi S_b, vi S_e, bool head, bool ceil, bool floor,
std::map<char,int>& nt2int, vi E_b, vi G_b)
{
    int F;
    *(G_b)=0;
    for(int j=0; j<S_e-S_b; ++j)
        for(int s=*(S_b+j)+1; s<=*(S_b+j+1); ++s)
        {
            if(ceil)
                *(G_b+s)=*(G_b+s-1)+((s==*(S_b+j)+1)?(*(tvf_b+j)):(*(tuf_b+j)));
            else
                *(G_b+s)=*(G_b+s-1)+((s==*(S_b+j)+1)?(*(vf_b+j)):(*(uf_b+j)));
            *(E_b+s)=std::numeric_limits<int>::min()/2;
        }
    for(int w=1; w<=o_e-o_b; ++w)
    {
        int hatG=*G_b;
        *E_b=*G_b=*G_b+((w==1 && head)?(*ve_b):(*ue_b));
        for(int j=0; j<S_e-S_b; ++j)
```

```
            for(int s=*(S_b+j)+1; s<=*(S_b+j+1); ++s)
            {
                if(floor && s==*(S_b+j+1))
                {
                    F=std::numeric_limits<int>::min()/2;
                    for(int s=*(S_b+j)+1; s<=*(S_b+j+1); ++s)
                        F=std::max(F,*(G_b+s-1)+*(tvf_b+j)+(*(S_b+j+1)-s)*(*(tuf_b+j)));
                }
                else
                {
                    if(s!=*(S_b+j)+1)
                        F=std::max(F+*(uf_b+j+w*row),*(G_b+s-1)+*(vf_b+j+w*row));
                    else
                        F=*(G_b+s-1)+*(vf_b+j+w*row);
                    if(ceil)
                        F=std::max(F,*(G_b+*(S_b+j))+*(tvf_b+j)+(s-*(S_b+j)-
1)*(*(tuf_b+j)));
                }
                *(E_b+s)=std::max(*(G_b+s)+*(ve_b+s),*(E_b+s)+*(ue_b+s));
                int Gp=*(G_b+s);
                *(G_b+s)=hatG+gamma(nt2int[*(x_b+s-1)],nt2int[*(o_b+w-1)],0);
                *(G_b+s)=std::max(*(G_b+s),*(E_b+s));
                *(G_b+s)=std::max(*(G_b+s),F);
                hatG=Gp;
            }
    }
}

void load_ref(std::string& ref_file, std::string& x, std::vector<int>& S, std::vector<int>&
left_exp, std::vector<int>& left_down, std::vector<int>& left_up, std::vector<int>& right_exp,
std::vector<int>& right_down, std::vector<int>& right_up)
{
    S.push_back(0);
    std::ifstream fin(ref_file);
    int le_tmp, ld_tmp, lu_tmp, re_tmp, rd_tmp, ru_tmp;
    std::string str_tmp;
    while(fin >> re_tmp >> rd_tmp >> ru_tmp >> str_tmp >> le_tmp >> ld_tmp >>
lu_tmp)
    {
        std::transform(str_tmp.begin(), str_tmp.end(), str_tmp.begin(), toupper);
        str_tmp.front()=tolower(str_tmp.front());
        str_tmp.back()=tolower(str_tmp.back());
        S.push_back(S.back()+str_tmp.size());
        x.append(str_tmp);
```

```
            left_exp.push_back(le_tmp);
            left_down.push_back(ld_tmp);
            left_up.push_back(lu_tmp);
            right_exp.push_back(re_tmp);
            right_down.push_back(rd_tmp);
            right_up.push_back(ru_tmp);
        }
        fin.close();
}

std::tuple<std::vector<std::string>,std::vector<int>,std::vector<double>>
load_read(std::vector<std::pair<std::vector<int>,std::string>>& sus_ord, int& so, int&
hasgot, int& max_len, int SEQ_BATCH, double per_thres, int total_read, std::ofstream& fout)
{
        hasgot=0;
        max_len=std::numeric_limits<int>::min();
        std::vector<std::string> os;
        std::vector<int> index;
        std::vector<double> num;
        while(so<sus_ord.size() &&
double(sus_ord[so].first.size())/double(total_read)>=per_thres)
        {
            fout << sus_ord[so].first.size() << '\t' << sus_ord[so].second << '\n';
            os.push_back(std::move(sus_ord[so].second));
            num.push_back(sus_ord[so].first.size());
            index.push_back(++so);
            ++hasgot;
            max_len=std::max(max_len,int(os.back().size()));
            if(hasgot==SEQ_BATCH)
                break;
        }
        return std::make_tuple(os,index,num);
}

std::vector<std::string> my_split(std::string line, std::string sc)
{
        std::vector<std::string> parts;
        std::string part;
        for(unsigned i=0; i<line.size(); ++i)
        {
            if(sc.find(line[i]) != std::string::npos)
            {
                parts.push_back(std::move(part));
                part.clear();
```

```
		}
		else
			part.push_back(line[i]);
	}
	parts.push_back(std::move(part));
	return parts;
}

Command_content command(int argc, char **argv)
{
	Command_content cc;

	for(int i=1; i<argc-1; ++i)
	{
		if(!strcmp(argv[i],"-files"))
			cc.files=my_split(argv[i+1],",");
		if(!strcmp(argv[i],"-disp_left"))
			cc.disp_left=my_split(argv[i+1],",");
		if(!strcmp(argv[i],"-disp_right"))
			cc.disp_right=my_split(argv[i+1],",");
		if(!strcmp(argv[i],"-disp_num"))
			cc.disp_num=my_split(argv[i+1],",");
		if(!strcmp(argv[i],"-DIVCON"))
			cc.DIVCON=my_split(argv[i+1],",");
		if(!strcmp(argv[i],"-SEQ_BATCH"))
			cc.SEQ_BATCH=my_split(argv[i+1],",");
		if(!strcmp(argv[i],"-ALIGN_MAX"))
			cc.ALIGN_MAX=my_split(argv[i+1],",");
		if(!strcmp(argv[i],"-s0"))
			cc.S0=my_split(argv[i+1],",");
		if(!strcmp(argv[i],"-s1"))
			cc.S1=my_split(argv[i+1],",");
		if(!strcmp(argv[i],"-u"))
			cc.u=my_split(argv[i+1],",");
		if(!strcmp(argv[i],"-v"))
			cc.v=my_split(argv[i+1],",");
		if(!strcmp(argv[i],"-references"))
			cc.references=my_split(argv[i+1],",");
		if(!strcmp(argv[i],"-alg_types"))
			cc.alg_types=my_split(argv[i+1],",");
		if(!strcmp(argv[i],"-per_thres"))
			cc.per_thres=my_split(argv[i+1],",");
		if(!strcmp(argv[i],"-ini_alpha"))
			cc.ini_alpha=my_split(argv[i+1],",");
```

```
		if(!strcmp(argv[i],"-ini_beta"))
			cc.ini_beta=my_split(argv[i+1],",");
		if(!strcmp(argv[i],"-ini_pi"))
			cc.ini_pi=my_split(argv[i+1],",");
		if(!strcmp(argv[i],"-MID_MAX"))
			cc.MID_MAX=my_split(argv[i+1],",");
		if(!strcmp(argv[i],"-thres"))
			cc.thres=my_split(argv[i+1],",");
		if(!strcmp(argv[i],"-mode"))
			cc.mode=my_split(argv[i+1],",");
		if(!strcmp(argv[i],"-THR_MAX"))
			cc.THR_MAX=my_split(argv[i+1],",");
	}

	if(cc.files.empty())
		getFiles(cc.files, cc.references);
	if(cc.disp_left.empty())
		cc.disp_left.push_back("20");
	if(cc.disp_right.empty())
		cc.disp_right.push_back("40");
	if(cc.disp_num.empty())
		cc.disp_num.push_back("20");
	if(cc.DIVCON.empty())
		cc.DIVCON.push_back("on");
	if(cc.SEQ_BATCH.empty())
		cc.SEQ_BATCH.push_back("300");
	if(cc.ALIGN_MAX.empty())
		cc.ALIGN_MAX.push_back("5");
	if(cc.S0.empty())
		cc.S0.push_back("-3");
	if(cc.S1.empty())
		cc.S1.push_back("1");
	if(cc.u.empty())
		cc.u.push_back("-2");
	if(cc.v.empty())
		cc.v.push_back("-5");
	if(cc.alg_types.empty())
		cc.alg_types.push_back("local_imbed");
	if(cc.per_thres.empty())
		cc.per_thres.push_back("0");
	if(cc.ini_alpha.empty())
		cc.ini_alpha.push_back("");
	if(cc.ini_beta.empty())
		cc.ini_beta.push_back("");
```

```
	if(cc.ini_pi.empty())
		cc.ini_pi.push_back("");
	if(cc.MID_MAX.empty())
		cc.MID_MAX.push_back("0");
	if(cc.thres.empty())
		cc.thres.push_back("0.000001");
	if(cc.mode.empty())
		cc.mode.push_back("nonoverlapping");
	if(cc.THR_MAX.empty())
		cc.THR_MAX.push_back(std::to_string(std::thread::hardware_concurrency()));

	cc.disp_left.resize(cc.files.size(),cc.disp_left.back());
	cc.disp_right.resize(cc.files.size(),cc.disp_right.back());
	cc.disp_num.resize(cc.files.size(),cc.disp_num.back());
	cc.DIVCON.resize(cc.files.size(),cc.DIVCON.back());
	cc.SEQ_BATCH.resize(cc.files.size(),cc.SEQ_BATCH.back());
	cc.ALIGN_MAX.resize(cc.files.size(),cc.ALIGN_MAX.back());
	cc.S0.resize(cc.files.size(),cc.S0.back());
	cc.S1.resize(cc.files.size(),cc.S1.back());
	cc.u.resize(cc.files.size(),cc.u.back());
	cc.v.resize(cc.files.size(),cc.v.back());
	cc.references.resize(cc.files.size(),cc.references.back());
	cc.alg_types.resize(cc.files.size(),cc.alg_types.back());
	cc.per_thres.resize(cc.files.size(),cc.per_thres.back());
	if(!cc.ini_alpha.empty())
		cc.ini_alpha.resize(cc.files.size(),cc.ini_alpha.back());
	if(!cc.ini_beta.empty())
		cc.ini_beta.resize(cc.files.size(),cc.ini_beta.back());
	if(!cc.ini_pi.empty())
		cc.ini_pi.resize(cc.files.size(),cc.ini_pi.back());
	cc.MID_MAX.resize(cc.files.size(),cc.MID_MAX.back());
	cc.thres.resize(cc.files.size(),cc.thres.back());
	cc.mode.resize(cc.files.size(),cc.mode.back());
	return cc;
}

void getFiles(std::vector<std::string> & files, std::vector<std::string> & references)
{
	const std::experimental::filesystem::path current_path(".");
	std::experimental::filesystem::directory_iterator end_itr; // default construction yields
past-the-end
	for ( std::experimental::filesystem::directory_iterator itr( current_path ); itr != end_itr;
++itr )
		if ( ! std::experimental::filesystem::is_directory(itr->status()) )
```

```
        {
            std::string str_tmp=itr->path().filename().string();
            if(std::find(references.begin(), references.end(), str_tmp) == references.end())
                files.push_back(str_tmp);
        }
    std::sort(files.begin(),files.end());
}

std::tuple<std::vector<int>,std::vector<int>,std::vector<std::string>,std::vector<int>,std::vect
or<int>,std::vector<std::string>,std::vector<int>,std::vector<int>,std::vector<std::string>>
search_left_right(std::pair<std::string,std::string>& alignment, std::vector<int>& S,
std::vector<int>& left_exp, std::vector<int>& right_exp, std::string& mode, std::string& x)
{
    std::vector<int> left, right, c_left, c_right, c_left_in, c_right_in;
    std::vector<int> right_S(S.begin(),S.end()-1);
    std::vector<int> left_S(S.begin()+1,S.end());
    std::tie(left,right)=extract_to_pair(alignment, left_S, right_S);
    std::tie(c_left,c_right)=normal_2_c(left, right, left_exp, right_exp, mode);
    std::tie(c_left_in,c_right_in)=extract_to_pair(alignment, c_left, c_right);
    std::vector<std::string> MID=get_mid(alignment, left, right);
    std::vector<std::string> c_MID_in=get_mid(alignment, c_left_in, c_right_in);
    std::vector<std::string> c_MID;
    for(int i=0; i<MID.size(); ++i)
    {
        if(i==0)
            c_MID.push_back(MID[i]+x.substr(right[i],c_right[i]-right[i]));
        else if(i==MID.size()-1)
            c_MID.push_back(x.substr(c_left[i-1],left[i-1]-c_left[i-1])+MID[i]);
        else
            c_MID.push_back(x.substr(c_left[i-1],left[i-1]-c_left[i-
1])+MID[i]+x.substr(right[i],c_right[i]-right[i]));
    }
    return std::make_tuple(left, right, MID, c_left, c_right, c_MID, c_left_in, c_right_in,
c_MID_in);
}


std::tuple<std::vector<int>,std::vector<int>>
extract_to_pair(std::pair<std::string,std::string>& alignment, std::vector<int>& left,
std::vector<int>& right)
{
    std::vector<int> left_ext, right_ext;
    int ref_cum=0, pair_now;
    bool search;
```

```
	if(ref_cum==right[0])
		search=true;
	else
		search=false;
	for(int i=0; i<alignment.first.size(); ++i)
	{
		if(alignment.first[i]!='-')
		{
			++ref_cum;
			if(alignment.second[i]!='-')
			{
				pair_now=ref_cum;
				if(search)
				{
					right_ext.push_back(pair_now-1);
					search=false;
				}
			}
			if(ref_cum==left[left_ext.size()])
			{
				if(search || left[left_ext.size()]==right[left_ext.size()])
				{
					right_ext.push_back((right[right_ext.size()]+left[left_ext.size()])/2);
					search=false;
					left_ext.push_back(right_ext.back());
				}
				else
					left_ext.push_back(pair_now);
			}
			if(right_ext.size()<right.size() && ref_cum==right[right_ext.size()])
				search=true;
			if(left_ext.size()==left.size())
				break;
		}
	}
	return std::make_tuple(left_ext,right_ext);
}

std::tuple<std::vector<int>,std::vector<int>> normal_2_c(std::vector<int>& left,
std::vector<int>& right, std::vector<int>& left_exp, std::vector<int>& right_exp, std::string&
mode)
{
	std::vector<int> c_left, c_right;
	if(mode=="nonoverlapping")
```

```
	{
		for(int i=0; i<left.size(); ++i)
		{
			c_left.push_back(std::min(left[i],left_exp[i]));
			c_right.push_back(std::max(right[i],right_exp[i]));
		}
	}
	else if(mode=="overlapping")
	{
		c_right.push_back(right.front());
		for(int i=0; i<left.size()-1; ++i)
		{
			c_left.push_back(std::min(left[i],left_exp[i]+std::max(right[i+1]-
right_exp[i+1],0)));
			c_right.push_back(std::max(right[i+1],right_exp[i+1]+std::min(left[i]-
left_exp[i],0)));
		}
		c_left.push_back(left.back());
	}
	for(int i=0; i<left.size(); ++i)
	{
		if(c_right[i]>c_left[i])
		{
			if(c_right[i]==right[i])
				c_left[i]=c_right[i];
			else if(c_left[i]==left[i])
				c_right[i]=c_left[i];
		}
	}
	return std::make_tuple(c_left,c_right);
}

std::vector<std::string> get_mid(std::pair<std::string,std::string>& alignment,
std::vector<int>& left, std::vector<int>& right)
{
	std::vector<std::string> MID(1,std::string());
	int ref_cum=0;
	bool insert=true;
	for(int i=0; i<alignment.first.size(); ++i)
	{
		if(alignment.first[i]!='-')
		{
			++ref_cum;
			if(MID.size()<=right.size() && ref_cum==right[MID.size()-1]+1)
```

```
				insert=false;
			}
			if(insert && alignment.second[i]!='-')
				MID.back().push_back(alignment.second[i]);
			if(MID.size()<=left.size() && ref_cum==left[MID.size()-1])
			{
				MID.push_back(std::string());
				insert=true;
			}
		}
		return MID;
}

void disp_sample(std::vector<Align>& aligns, std::vector<int>& left_exp, std::vector<int>&
right_exp, int disp_left, int disp_right, int disp_num, std::string& x, std::string& mode,
std::string& file, int total_read)
{
		std::vector<JuncIndel> juncindels;
		for(int i=0; i<aligns.size(); ++i)
		{
			if(i!=0 && aligns[i].index==aligns[i-1].index) continue;
			juncindels.push_back(JuncIndel());
			juncindels.back().num=aligns[i].num;
			std::vector<int> c_left_in_alg, c_right_in_alg;
			int ref_cum=0;
			while(c_left_in_alg.size()<aligns[i].c_left_in.size() &&
ref_cum==aligns[i].c_left_in[c_left_in_alg.size()])
				c_left_in_alg.push_back(0);
			for(int j=0; j<aligns[i].alignment.first.size(); ++j)
			{
				if(aligns[i].alignment.first[j]!='-')
				{
					++ref_cum;
					while(c_left_in_alg.size()<aligns[i].c_left_in.size() &&
ref_cum==aligns[i].c_left_in[c_left_in_alg.size()])
						c_left_in_alg.push_back(j+1);
					while(c_right_in_alg.size()<aligns[i].c_right_in.size() &&
ref_cum==aligns[i].c_right_in[c_right_in_alg.size()]+1)
						c_right_in_alg.push_back(j);
					if(c_left_in_alg.size()==aligns[i].c_left_in.size() &&
c_right_in_alg.size()==aligns[i].c_right_in.size())
						break;
				}
			}
```

```
		for(int j=0; j<left_exp.size()-1; ++j)
		{
			int ocp_left=std::min(std::max(aligns[i].c_left_in[j]-left_exp[j],-
disp_left),disp_right);
			int ocp_right=std::max(std::min(aligns[i].c_right_in[j+1]-
right_exp[j+1],disp_right),-disp_left);
			if(ocp_left>ocp_right)
				ocp_right=ocp_left;
			std::string del(ocp_right-ocp_left,'-');
			std::string
ins=aligns[i].c_MID_in[j+1].substr(0,std::min(int(aligns[i].c_MID_in[j+1].size()),disp_right-
ocp_right));
			ocp_right+=ins.size();
			int tmp=std::max(0,c_left_in_alg[j]-ocp_left-disp_left);
			std::string lep=aligns[i].alignment.second.substr(tmp,c_left_in_alg[j]-tmp);
			std::vector<int> lep_indel;
			for(int k=tmp; k<c_left_in_alg[j]; ++k)
			{
				if(aligns[i].alignment.second[k]!='-' && aligns[i].alignment.first[k]=='-')
					lep_indel.push_back(1);
				else if(aligns[i].alignment.second[k]!='-' &&
aligns[i].alignment.first[k]!=aligns[i].alignment.second[k])
					lep_indel.push_back(2);
				else
					lep_indel.push_back(0);
			}
			tmp=ocp_left+disp_left-lep.size();
			lep=std::string(tmp,'-')+lep;
			lep_indel.insert (lep_indel.begin(),tmp,0);
			tmp=std::min(c_right_in_alg[j+1]+disp_right-
ocp_right,int(aligns[i].alignment.second.size()));
			std::string rip=aligns[i].alignment.second.substr(c_right_in_alg[j+1],tmp-
c_right_in_alg[j+1]);
			std::vector<int> rip_indel;
			for(int k=c_right_in_alg[j+1]; k<tmp; ++k)
			{
				if(aligns[i].alignment.second[k]!='-' && aligns[i].alignment.first[k]=='-')
					rip_indel.push_back(1);
				else if(aligns[i].alignment.second[k]!='-' &&
aligns[i].alignment.first[k]!=aligns[i].alignment.second[k])
					rip_indel.push_back(2);
				else
					rip_indel.push_back(0);
			}
```

```
            rip.resize(disp_right-ocp_right,'-');
            rip_indel.resize(disp_right-ocp_right,0);
            juncindels.back().insert.push_back(std::move(lep_indel));
            juncindels.back().insert.push_back(std::vector<int>(del.size(),0));
            juncindels.back().insert.push_back(std::vector<int>(ins.size(),1));
            juncindels.back().insert.push_back(std::move(rip_indel));
            juncindels.back().ligation_indel.push_back(std::move(lep));
            juncindels.back().ligation_indel.push_back(std::move(del));
            juncindels.back().ligation_indel.push_back(std::move(ins));
            juncindels.back().ligation_indel.push_back(std::move(rip));
        }
        if(juncindels.size()==disp_num) break;
    }

    for(int i=0; i<left_exp.size()-1; ++i)
    {
        std::ofstream fout_heat(file+"."+mode+std::to_string(i+1)+".heat"),
fout_letter(file+"."+mode+std::to_string(i+1)+".letter"),
fout_line(file+"."+mode+std::to_string(i+1)+".line"),
fout_word(file+"."+mode+std::to_string(i+1)+".word"),
fout_junc(file+"."+mode+std::to_string(i+1)+".junc");
        int tmp=std::max(0,left_exp[i]-disp_left);
        std::string ref=std::string(tmp-left_exp[i]+disp_left,'-')+x.substr(tmp,left_exp[i]-
tmp)+x.substr(right_exp[i+1],disp_right);
        ref.resize(disp_right+disp_left,'-');
        enum {BLACK=0, RED=65536*255 + 256*0 + 0, BLUE=65536*0 + 256*0 + 255};
        std::map<char,std::array<int,3>>
char_map_RGB{{'A',{0,255,0}},{'T',{128,0,128}},{'G',{255,255,0}},{'C',{0,255,255}},{'N',{255,255,25
5}},{'-',{255,255,255}}};
        for(int j=0; j<ref.size(); ++j)
        {
            ref[j]=toupper(ref[j]);
            std::array<int,3> tmp=char_map_RGB[ref[j]];
            fout_heat << j+0.5 << '\t' << 0.5 << '\t' << tmp[0] << '\t' << tmp[1] << '\t'
<< tmp[2] << '\n';
            fout_letter << j+0.5 << '\t' << 0.5 << '\t' << ref[j] << '\t' << BLACK << '\n';
        }
        fout_word     << 2 << '\t' << juncindels.size()+1.5 << '\t' << "Deletions" << '\t'
<< BLACK << '\n'
                      << 5 << '\t' << juncindels.size()+2.5 << '\t' << "Insertions" << '\t'
<< BLACK << '\n'
                      << 5 << '\t' << juncindels.size()+3.5 << '\t' << "Substitutions" <<
'\t' << BLACK << '\n'
```

```
                    << 5 << '\t' << juncindels.size()+4.5 << '\t' << "\"Ligation indel\""
<< '\t' << BLACK << '\n'
                    << 1 << '\t' << juncindels.size()+5.5 << '\t' << "Ligation" << '\t' <<
BLACK << '\n'
                    << ref.size()+1 << '\t' << 0.5 << '\t' << "Reference" << '\t' <<
BLACK << '\n';
        fout_letter << 0.5 << '\t' << juncindels.size()+1.5 << '\t' << "-" << '\t' << BLACK
<< '\n'
                    << 0.5 << '\t' << juncindels.size()+2.5 << '\t' << "A" << '\t' << RED
<< '\n'
                    << 1.5 << '\t' << juncindels.size()+2.5 << '\t' << "T" << '\t' << RED
<< '\n'
                    << 2.5 << '\t' << juncindels.size()+2.5 << '\t' << "G" << '\t' << RED
<< '\n'
                    << 3.5 << '\t' << juncindels.size()+2.5 << '\t' << "C" << '\t' << RED
<< '\n'
                    << 0.5 << '\t' << juncindels.size()+3.5 << '\t' << "A" << '\t' <<
BLUE << '\n'
                    << 1.5 << '\t' << juncindels.size()+3.5 << '\t' << "T" << '\t' <<
BLUE << '\n'
                    << 2.5 << '\t' << juncindels.size()+3.5 << '\t' << "G" << '\t' <<
BLUE << '\n'
                    << 3.5 << '\t' << juncindels.size()+3.5 << '\t' << "C" << '\t' <<
BLUE << '\n';
        fout_junc   << disp_left << '\t' << 0 << '\t' << 0 << '\t' << 1 << '\t' << BLACK
<< '\n'
                    << 0 << '\t' << juncindels.size()+5 << '\t' << 0 << '\t' << 1 << '\t'
<< BLACK << '\n';
        fout_line   << 0 << '\t' << juncindels.size()+4 << '\t' << 0 << '\t' << 1 << '\t' <<
BLACK << '\n'
                    << 0 << '\t' << juncindels.size()+4 << '\t' << 4 << '\t' << 0 << '\t'
<< BLACK << '\n'
                    << 0 << '\t' << juncindels.size()+5 << '\t' << 4 << '\t' << 0 << '\t'
<< BLACK << '\n'
                    << 4 << '\t' << juncindels.size()+4 << '\t' << 0 << '\t' << 1 << '\t'
<< BLACK << '\n';

        std::streamsize ss = fout_word.precision();
        for(int sn=0; sn<juncindels.size(); ++sn)
        {
            fout_word << ref.size()+1 << '\t' << sn+1.5 << '\t' <<
setiosflags(std::ios::fixed) << std::setprecision(2) <<
juncindels[sn].num*100/double(total_read) << "%" << resetiosflags(std::ios::fixed) <<
std::setprecision(ss) <<'\t' << BLACK << '\n';
```

```
			fout_word << ref.size()+7 << '\t' << sn+1.5 << '\t' << "\"" << juncindels[sn].num << " reads\"" << '\t' << BLACK << '\n';
			fout_line << juncindels[sn].ligation_indel[i*4+0].size() << '\t' << sn+1 << '\t' << 0 << '\t' << 1 << '\t' << BLACK << '\n';
			fout_line << juncindels[sn].ligation_indel[i*4+0].size() << '\t' << sn+1 << '\t' << juncindels[sn].ligation_indel[i*4+1].size()+juncindels[sn].ligation_indel[i*4+2].size() << '\t' << 0 << '\t' << BLACK << '\n';
			fout_line << juncindels[sn].ligation_indel[i*4+0].size() << '\t' << sn+2 << '\t' << juncindels[sn].ligation_indel[i*4+1].size()+juncindels[sn].ligation_indel[i*4+2].size() << '\t' << 0 << '\t' << BLACK << '\n';
			fout_line << juncindels[sn].ligation_indel[i*4+0].size()+juncindels[sn].ligation_indel[i*4+1].size()+juncindels[sn].ligation_indel[i*4+2].size() << '\t' << sn+1 << '\t' << 0 << '\t' << 1 << '\t' << BLACK << '\n';

			int pos=0;
			for(int sg=0; sg<4; sg++)
			{
				for(int j=0; j<juncindels[sn].ligation_indel[i*4+sg].size(); ++j)
				{
					juncindels[sn].ligation_indel[i*4+sg][j]=toupper(juncindels[sn].ligation_indel[i*4+sg][j]);
					std::array<int,3> tmp=char_map_RGB[juncindels[sn].ligation_indel[i*4+sg][j]];
					fout_heat << pos+0.5 << '\t' << sn+1.5 << '\t' << tmp[0] << '\t' << tmp[1] << '\t' << tmp[2] << '\n';
					fout_letter << pos+0.5 << '\t' << sn+1.5 << '\t' << juncindels[sn].ligation_indel[i*4+sg][j] << '\t';
					if(juncindels[sn].insert[i*4+sg][j]==1)
						fout_letter << RED;
					else if(juncindels[sn].insert[i*4+sg][j]==2)
						fout_letter << BLUE;
					else
						fout_letter << BLACK;
					fout_letter << '\n';
					++pos;
				}
			}
		}
		fout_heat.close();
		fout_letter.close();
		fout_line.close();
		fout_word.close();
		fout_junc.close();
```

```
		std::ofstream fout_gp(file+"."+mode+std::to_string(i+1)+".gp");
		fout_gp << "set terminal postscript eps color font \"Arial, 12\" linewidth 1.0\n";
		fout_gp << "set output \""+file+"."+mode+std::to_string(i+1)+".eps\"\n";
		fout_gp << "set size 1.0,1.0\n";
		fout_gp << "unset border\n";
		fout_gp << "set encoding utf8\n";
		fout_gp << "set xrange [0:"+std::to_string(ref.size()+15)+"] noextend\n";
		fout_gp << "set yrange ["+std::to_string(juncindels.size()+6)+":0] noextend\n";
		fout_gp << "unset xlabel\n";
		fout_gp << "unset ylabel\n";
		fout_gp << "unset xtics\n";
		fout_gp << "unset ytics\n";
		fout_gp << "plot \'"+file+"."+mode+std::to_string(i+1)+".heat\' with rgbimage
notitle, \'"+file+"."+mode+std::to_string(i+1)+".letter\' with labels tc rgb variable font
\"Courier, 12\" notitle, \'"+file+"."+mode+std::to_string(i+1)+".line\' with vectors lc rgb
variable lw 2 nohead notitle, \'"+file+"."+mode+std::to_string(i+1)+".junc\' with vectors lc
rgb variable lw 2 dt (2,4) nohead notitle, \'"+file+"."+mode+std::to_string(i+1)+".word\' with
labels tc rgb variable left notitle\n";
		fout_gp.close();
	}
}

void indel_label_fun(std::vector<Align>& aligns,
std::vector<std::pair<std::vector<int>,std::string>>& sus_ord, int total_read,
std::vector<int>& left_exp, std::vector<int>& right_exp, std::string& file)
{
	std::vector<std::vector<std::array<int,3>>> LPRPILs(total_read);
	std::array<int,3> LPRPIL;
	for(auto & align : aligns)
	{
		for(int i=1; i<right_exp.size(); ++i)
		{

			LPRPIL[0]=align.left[i-1]-left_exp[i-1];
			LPRPIL[1]=align.right[i]-right_exp[i];
			LPRPIL[2]=align.MID[i].size();
			for(auto j : sus_ord[align.index-1].first)
				LPRPILs[j].push_back(LPRPIL);
		}
	}
	std::ofstream fout(file+".indel");
	for(auto & LPRPIL_row : LPRPILs)
	{
```

```
		if(LPRPIL_row.empty())
			fout << '\n';
		for(int i=0; i<LPRPIL_row.size(); ++i)
		{
			fout << LPRPIL_row[i][0] << '\t' << LPRPIL_row[i][1] << '\t' <<
LPRPIL_row[i][2];
			if(i!=LPRPIL_row.size()-1)
				fout << '\t';
			else
				fout << '\n';
		}
	}
	fout.close();
}

void div_num_fun(std::vector<Align>::iterator first, std::vector<Align>::iterator last)
{
	if (first==last)
		return;
	std::vector<Align>::iterator tracer = first;
	double index_num=1;
	while (++first != last)
	{
		if((first-1)->index!=first->index)
		{
			for(;tracer<first;++tracer)
			{
				tracer->num/=index_num;
			}
			index_num=1;
		}
		else
			index_num+=1;
	}
	for(;tracer<first;++tracer)
	{
		tracer->num/=index_num;
	}
}

void EM_predict(std::vector<Align>& aligns, std::string& x, std::vector<int>& left_exp,
std::vector<int>& left_down, std::vector<int>& left_up, std::vector<int>& right_exp,
std::vector<int>& right_down, std::vector<int>& right_up, int MID_MAX, std::string&
ini_alpha, std::string& ini_beta, std::string& ini_pi, double thres, std::string& file)
```

```
{
	div_num_fun(aligns.begin(), aligns.end());

	std::vector<std::string> pos_2_MID;
		std::string ind_tmp("NACGT");
		for(int k=0;k<=MID_MAX;++k)
		{
			for(int w=0;w<std::pow(5,k);++w)
			{
				std::string MID;
				for(int i=0;i<k;i++)
				{
					MID+=ind_tmp[w%5];
					w/=5;
				}
				pos_2_MID.push_back(std::move(MID));
			}
		}


std::vector<std::map<std::tuple<int,int,std::string>,std::pair<double,std::vector<double*>>,
MY_LESS>> tuple_2_numpoi;
	std::vector<TD_array<double>> N_mats;
	for(int i=1; i<right_down.size(); ++i)
	{
		int n_row=left_up[i-1]-left_down[i-1]+1, n_col=right_up[i]-right_down[i]+1,
n_slice=int((1-pow(5,MID_MAX+1))/(1-5)+0.5);
		N_mats.push_back(TD_array<double>(n_row, n_col, n_slice));

tuple_2_numpoi.push_back(std::map<std::tuple<int,int,std::string>,std::pair<double,std::vect
or<double*>>,MY_LESS>(MY_LESS(x)));
		for(int r=0; r<n_row; ++r)
			for(int c=0; c<n_col; ++c)
				for(int s=0; s<n_slice; ++s)
				{
					std::tuple<int,int,std::string> tmp=std::make_tuple(r+left_down[i-
1],c+right_down[i],pos_2_MID[s]);
					tuple_2_numpoi.back()[tmp].first=0;

tuple_2_numpoi.back()[tmp].second.push_back(&N_mats.back()(r,c,s));
				}

		for(auto& align : aligns)
		{
```

```
            std::tuple<int,int,std::string> tmp=std::make_tuple(align.left[i-1],align.right[i],align.MID[i]);
            if(tuple_2_numpoi.back().count(tmp)>0)
                tuple_2_numpoi.back()[tmp].first+=align.num;
        }

    }
    std::string pre_str;
    std::experimental::filesystem::path pathObj;
    pathObj=ini_alpha;
    pre_str.append(pathObj.filename().string());
    pathObj=ini_beta;
    if(!pre_str.empty() && !pathObj.filename().string().empty())
        pre_str.append("_");
    pre_str.append(pathObj.filename().string());
    pathObj=ini_pi;
    if(!pre_str.empty() && !pathObj.filename().string().empty())
        pre_str.append("_");
    pre_str.append(pathObj.filename().string());
    if(!pre_str.empty())
        pre_str.append("_");
    std::vector<std::vector<double>> alphas=initial_EM(ini_alpha, 0, N_mats);
    std::vector<std::vector<double>> betas=initial_EM(ini_beta, 1, N_mats);
    std::vector<std::vector<double>> pis=initial_EM(ini_pi, 2, N_mats);
    for(int i=0; i<left_down.size()-1; ++i)
    {
        double sum_tmp=0;
        for(auto & pair : tuple_2_numpoi[i])
            sum_tmp+=pair.second.first;
        if(sum_tmp==0)
        {
            std::fill(alphas[i].begin(),alphas[i].end(),0.0);
            std::fill(betas[i].begin(),betas[i].end(),0.0);
            std::fill(pis[i].begin(),pis[i].end(),0.0);
            N_mats[i].fill(0.0);
        }
        else
        {
            double tol=std::numeric_limits<double>::max();
            while(tol>thres)
            {
                // E-step
                for(int r=0;r<alphas[i].size();++r)
                    for(int c=0;c<betas[i].size();++c)
```

```
                    for(int s=0;s<pis[i].size();++s)
                        N_mats[i](r,c,s)=alphas[i][r]*betas[i][c]*pis[i][s];
                for(auto & pair : tuple_2_numpoi[i])
                {
                    double accum=0;
                    for(auto & add : pair.second.second)
                        accum+=*add;
                    if(accum>0)
                        for(auto & add : pair.second.second)
                            *add=pair.second.first*(*add)/accum;
                    else
                        for(auto & add : pair.second.second)
                            *add=pair.second.first/pair.second.second.size();
                }
                // M-step
                std::vector<double> alpha_old=alphas[i], beta_old=betas[i], pi_old=pis[i];
                std::tie(alphas[i],betas[i],pis[i])=N_mats[i].boundary_sum();
                for(auto & comp : alphas[i]) comp/=sum_tmp;
                for(auto & comp : betas[i]) comp/=sum_tmp;
                for(auto & comp : pis[i]) comp/=sum_tmp;

                tol=0;
                for(int r=0; r<alpha_old.size(); ++r)
                {
                    double tmp=std::abs(alpha_old[r]-alphas[i][r]);
                    if(tmp>tol) tol=tmp;
                }
                for(int c=0; c<beta_old.size(); ++c)
                {
                    double tmp=std::abs(beta_old[c]-betas[i][c]);
                    if(tmp>tol) tol=tmp;
                }
                for(int s; s<pi_old.size(); ++s)
                {
                    double tmp=std::abs(pi_old[s]-pis[i][s]);
                    if(tmp>tol) tol=tmp;
                }
            }
        }
        for(int r=0;r<alphas[i].size();++r)
            for(int c=0;c<betas[i].size();++c)
                for(int s=0;s<pis[i].size();++s)
                    N_mats[i](r,c,s)=alphas[i][r]*betas[i][c]*pis[i][s];
        std::ofstream fout(pre_str+file+std::to_string(i+1)+".box");
```

```
            for(auto & pair : tuple_2_numpoi[i])
            {
                  double accum=0;
                  for(auto & add : pair.second.second)
                        accum+=*add;
                  fout << pair.second.first << '\t' << accum*sum_tmp << '\t' <<
pair.second.second.size() << '\n';
            }
            fout.close();
      }
      std::ofstream fout;
      fout.open(pre_str+file+".alpha");
      for(int i=0; i<left_exp.size()-1; ++i)
            for(int j=left_down[i]; j<=left_up[i]; ++j)
                  fout << j-left_exp[i] << '\t' << alphas[i][j-left_down[i]] << '\n';
      fout.close();
      fout.open(pre_str+file+".beta");
      for(int i=1; i<right_exp.size(); ++i)
            for(int j=right_down[i]; j<=right_up[i]; ++j)
                  fout << j-right_exp[i] << '\t' << betas[i-1][j-right_down[i]] << '\n';
      fout.close();
      fout.open(pre_str+file+".pi");
      for(int i=0; i<left_exp.size()-1; ++i)
            for(int j=0; j<pos_2_MID.size(); ++j)
                  fout << pos_2_MID[j] << '\t' << pis[i][j] << '\n';
      fout.close();
}

std::vector<std::vector<double>> initial_EM(std::string& ini, int dim,
std::vector<TD_array<double>>& N_mats)
{
      std::vector<std::vector<double>> vecs;
      if(ini.empty())
            for(auto & N_mat : N_mats)
            {
                  int tmp=N_mat.size(dim);
                  vecs.push_back(std::vector<double>(tmp,1.0/tmp));
            }
      else
      {
            std::ifstream fin(ini);
            for(auto & N_mat : N_mats)
            {
                  int tmp=N_mat.size(dim);
```

```
            vecs.push_back(std::vector<double>());
            std::string line;
            for(int rein=0; rein<tmp; ++rein)
            {
                getline(fin,line);
                vecs.back().push_back(stod(my_split(line," \t\n\v\f\r")[1]));
            }
        }
        fin.close();
    }
    return vecs;
}
```

### **Supplementary Note S5.** MATLAB script

```
function longest_common_substring_cumulative(n, M, p, outpath)
b = M - 2 * log(n) ./ log(1/p);
proupper = (1+p) ./ (1-p) .* (1 - 2 ./ n .* log(n) ./ log(1/p)) .^ (-2) .* p .^ (-b-1);
proupper = min(proupper, 1);
prolower = 1 - p .^ (b + 1);
prolower = max(prolower, 0);
writetable(table(n(:), prolower(:), proupper(:), 'VariableNames', {'length',
'lower','upper'}),fullfile(outpath,sprintf('%d.mmej.table',M)), 'FileType', 'text', 'Delimiter', '\t');
h_fig=figure('Position',[0 0 1800 900], 'Visible', 'off');
ax=axes(h_fig,'Position',[0.05,0.05,0.9,0.9]);
hold(ax,'on');
plot(n, proupper, '.-k', n, prolower, '.-r');
legend(ax, 'upper', 'lower');
ax.XLim = [min(n), max(n)];
ax.XTick = ax.XLim;
ax.YTick = ax.YLim;
axes_regular_setting(ax);
ax.YTickLabels = compose('%g',ax.YTick);
xlabel(ax, 'length');
ylabel(ax, 'bounds of probability of no mmej');
print('-painters', h_fig, fullfile(outpath, sprintf('%d.nommej.png',M)), '-dpng');
print('-painters', h_fig, fullfile(outpath, sprintf('%d.nommej.eps',M)), '-depsc');

h_fig=figure('Position',[0 0 1800 900], 'Visible', 'off');
ax=axes(h_fig,'Position',[0.05,0.05,0.9,0.9]);
hold(ax,'on');
plot(n, 1-proupper, '.-k');
```

```
legend(ax, 'lower');
ax.XLim = [min(n), max(n)];
ax.XTick = ax.XLim;
ax.YTick = ax.YLim;
axes_regular_setting(ax);
ax.YTickLabels = compose('%g',ax.YTick);
xlabel(ax, 'length');
ylabel(ax, 'lower bound of probability of mmej');
print('-painters', h_fig, fullfile(outpath, sprintf('%d.mmej.png',M)), '-dpng');
print('-painters', h_fig, fullfile(outpath, sprintf('%d.mmej.eps',M)), '-depsc');
end
```
